# Single-cell sequencing characterizes intratumoral profile of immune cells in old mice

**DOI:** 10.1101/2021.02.23.432366

**Authors:** Cangang Zhang, Lei Lei, Xiaofeng Yang, Huiqiang Zheng, Yanhong Su, Anjun Jiao, Xin Wang, Haiyan Liu, Yujing Zou, Lin Shi, Xiaobo Zhou, Chenming Sun, Lianjun Zhang, Yuzhu Hou, Zhengtao Xiao, Baojun Zhang

## Abstract

It has long been thought that aging is a major risk factor for cancer incidence. However, accumulating evidence indicates increased resistance of old animals to tumor growth. A systematic understanding of how old individuals defend against tumor invasion is currently lacking. Here we investigated the differences of age-associated alterations in tumor-infiltrating immune cells between young and old mice using single-cell RNA analysis. Our results showed that a higher proportion of cytotoxic CD8^+^ T cells, nTregs, cDC, and M1-type macrophages, while a higher percentage of exhausted CD8^+^ T cells, iTregs, pDC, and M2-type macrophages were found in young mice. Importantly, TCR diversity analysis showed top 10 TCR clones consisted primarily of exhausted CD8^+^ T cells in young mice whereas tip clones were predominantly cytotoxic CD8^+^ T cells in old mice. Consistently, trajectory inference demonstrated that CD8^+^ T cells preferentially differentiated into cytotoxic cells in old mice in contrast to exhausted cells in young mice. Meanwhile, we confirmed the main distinctions between young and old mice by flow cytometry. Collectively, our data revealed that a significantly higher proportion of effector immune cells in old mice defend against tumor progression, providing a framework for the immunotherapy of elderly patients with tumors.

## Introduction

Aging leads to the inevitable time-dependent decline in organ function and is a major risk factor for cancer. Relevant mechanisms of higher cancer incidence in aging individuals include genomic instability, epigenetic changes, loss of proteostasis, and declining immune surveillance (1). Individuals with an aging immune system are more susceptible to infection, and experience reduced vaccine effectiveness, higher incidence of cancer and autoimmune diseases (2, 3). More than 50% of cancers and approximately 70% of cancer-related deaths occur in patients older than 65 (4). Interestingly, many clinical and preclinical studies indicated that tumor in young patients or animals grow more aggressively than old counterparts (5–7). Furthermore, adoptive transfer of bone marrow or spleen cells from old mice has been shown to reduce the aggressiveness of tumor growth in young mice (5), paradoxical to the concept that cancer is defined as a disease of aging.

Over the past decade, immunotherapies that modulate the immune microenvironment to target and eliminate tumor cells have greatly changed the treatment approaches of cancer (8). The most common treatments that have proven clinical efficacy across a broad range of cancers are immune checkpoint inhibitors against programmed cell death 1(PD1), PDL1, and cytotoxic T lymphocyte-associated antigen 4 (CTLA4), both of which target T cells to restore their anti-tumor capacity. Immune checkpoint inhibitors significantly improve overall survival(OS) in both young and elder patients, but the magnitude of benefit is age-variable(1). Several clinical studies showed that elderly patients could benefit more from immunotherapies than young patients (9, 10) while a similar phenomenon is observed in mice implanted with melanoma tumors (11, 12). Therefore, it is of great significance to explore the immune system of elderly cancer patients and to further improve immunotherapy for the elderly.

The immune system plays a vital role in recognizing tumor cells and inhibiting the growth of malignant tissues (13). Several studies attempted to reveal the mechanisms of delayed tumor progression in the elderly from the perspective of anti-tumor immunity. It has been shown that an accumulation of 4-1BBL^+^ B cells in the elderly generated GranzymeB^+^ CD8^+^ T cells controls tumor growth (14), and old CD8^+^ T cells demonstrate increased adhesion and thus more easily infiltrated tumors through high expression of integrin α 4 (12). However, tumor-infiltrating immune cells form a complex mixture and the dense cross-talk of different types of immune cells plays a critical role in tumor immune surveillance (15). It is difficult to comprehensively characterize the immune profile to fully understand the mechanisms of delayed tumor development and the preferable outcome of immunotherapies in the elderly with conventional immunology technologies. As a revolutionary technology, single-cell RNA sequencing (single-cell RNA seq) allows for the ability to characterize cell types at the single-cell level and accurately define their various immune functions in the complex tumor microenvironment(TME) (16). Accumulating studies using single-cell RNA seq to explore tumor-infiltrating immune cells have been continuously reported (17–21). Besides, Single-cell RNA seq provides a powerful tool to define the T cell receptor (TCR) sequence and dominant clones recognizing tumor antigens (22).

In the current study, we utilized single-cell RNA seq to decipher the transcriptomic landscape of immune cells in B16 melanoma between young and old mice, and uncovered differences in the immune cell proportions and their functional features in young and old mouse tumors. We showed that intratumoral immune cells consist of more effector subsets including cytotoxic CD8^+^ T cells, cDC, and M1-type macrophages in old mice, while immunesuppressive subsets including exhausted CD8^+^ T cells, iTregs, pDC, and M2-type macrophages are primarily found in young mice. Importantly, TCR diversity analysis demonstrated that a significant proportion of cytotoxic CD8^+^ T cells were directly differentiated from effector memory like CD8^+^ T cells (CD8^+^ EM_like T) in the TME of old mice. Overall, our data demonstrated that immune cells in the TME of old mice hinder tumor growth, providing critical findings that can facilitate understanding of cancer development and immunotherapy efficacy in elderly patients.

## Materials and Methods

### Mice and reagents

6 to 8 week-old and 20 to 22 month-old female C57BL/6 mice were used in this study. All the mice were obtained from the Laboratory Animal Center of Xi’an Jiaotong University and were housed in the specific pathogen-free animal facility. All animal procedures were performed with the approval of the Animal Care Committee of Xi’an Jiaotong University and conformed to the Guide for the Care and Use of Laboratory Animals published by the US National Institutes of Health.

The antibodies and intracellular staining kit used were as follows: APC/Cy7 anti-mouse Cd4 (clone GK1.5), PE/Cy5 anti-mouse Cd4 (clone GK1.5), APC/Cy7 anti-mouse Cd8a (clone 53-6.7), Pacific BlueTM anti-mouse Cd8a (clone 53-6.7), PE/Cy7 anti-mouse/human Cd44 (clone IM7), PE/Cy5 anti-mouse Cd25 (clone PC61), PE/Cy5 anti-mouse Cd127(IL-7Rα) (clone A7R34), FITC anti-mouse CD11c (clone N418), FITC anti-mouse/human Cd11b (clone M1/70), PE/Cy7 anti-mouse I-A/I-E (clone M5/114.15.2), APC/Cy7 anti-mouse Cd19 (clone 6D5), Pacific BlueTM anti-mouse/human Cd45R/B220 (clone RA3-6B2), FITC anti-mouse Cd45.2 (clone 104), PE anti-mouse Cd45.2 (clone 104), PE/Cy7 anti-mouse IL-10 (clone JES5-16E3), Pacific BlueTM anti-mouse FOXP3 (clone MF-14), APC/Cy7 anti-mouse IFNγ (clone XMG1.2), APC anti-mouse LAP(TGF-β) (clone TW7-16B4), APC anti-mouse Cd223(LAG-3) (clone C9B7W), and Fixation Buffer(Cat # 420801) and Intracellular Staining Perm Wash Buffer(Cat # 421002). All above-mentioned reagents were purchased from BioLegend (San Diego, CA, USA). FITC anti-Mo Granzyme B (clone NGZB), and Transcription Factor Fixation/Permeabilization Concentrate and Diluent were purchased from eBioscience.

### Tumor model and preparation of cell suspensions

Two groups of mice were injected subcutaneously with 2×10^5^ B16F10 cells per mouse. After 5 days, tumor tissues were excised and weighed at indicated time points (every two days). Tumor volumes were determined by caliper measurement using the formula V = (length × width^2^) / 2. Freshly isolated tumor tissues were minced into approximately 1 mm^3^ cubic pieces and digested using 0.1% collagenase IV (LS004186, Worthington), 0.002% DNAse I (D8071, Solarbio), and 0.01% hyaluronidase (H3506-1G, SIGMA), then incubated on a rocker at 37°C for 40-50 min. The digested cells were filtered through a 70 μm cell strainer and washed twice with cold FACS buffer. The remaining cells were stained with CD45.2-APC (Clone 104; BioLegend) and 7AAD (Part 76332; Lot B226294 Biolegend) for 30 min at 4 °C, then washed and suspended in FACS buffer for flow cytometric sorting using FACS Aria II Cell Sorter (BD Biosciences). Sorted CD45.2^+^7AAD^-^ cells with a purity greater than 95% and viability higher than 90% were used for single-cell RNA sequencing.

### Single-cell RNA-seq and VDJ sequencing

The single-cell library preparation was constructed using 10X Chromium Single Cell V(D)J V2 Reagent Kits according to the manufacturer’s protocol. Briefly, single-cell suspensions with a concentration of 1000 cells/ul were loaded on the 10X genomics chromium controller single-cell instrument. Reverse transcription reagents, barcoded gel beads, and partitioning oil were mixed with the cells for generating single-cell gel beads in emulsions (GEM). After reverse transcription reaction, the GEMs were broken. The barcoded, full-length cDNA was amplified and purified to build V(D)J enriched TCR library and 5’ gene expression library. The mouse T Cell V (D) J Enrichment Kit was used to isolate and enrich for the V (D) J sequence. Finally, the constructed libraries were sequenced on the Illumina NovaSeq 6000 platform with NovaSeq 6000 S4 Reagent Kit (300 cycles).

### Single-cell RNA-seq data processing

Cell Ranger (v3.0.2,https://support.10xgenomics.com/) was used to process single-cell sequencing data and generate the matrix data containing gene counts for each cell per sample. Briefly, raw base call files from Illumina sequencers were first demultiplexed into FASTQ files with the cellranger mkfastq pipeline. Then, the splicing-aware aligner STAR was used to align FASTQs files to the mouse reference genome (mm10). The aligned reads were further counted using the cellranger count pipeline. Finally, the gene expression matrixes of all samples were imported into Seurat v3(23) and merged for subsequent analyses. The following filtering steps were carried out to exclude low-quality cells: cells with fewer than 200 and more than 3000 detected genes were discarded; cells with a high fraction of mitochondrial genes (>10%) are removed. As a result, a total of 4606 cells (young mice) and 5375 cells (old mice) with 973 informative genes were included in the analyses.

### Clustering of single cells and cell-type annotation

The gene expression data were log-normalized and scaled with default parameters. The top 3,000 most variable genes identified by Seurat function “FindVariableFeatures” were used for the principal component analysis (PCA). The first 50 principal components (PCs) selected based on the ElbowPlot were used for clustering analyses. Cell clusters were identified using FindClusters functions implemented in Seurat with default parameters and resolution parameter as 1.2. The t-SNE and UMAP were used to visualize the clustering results with default parameters and learning rate setting to 1.12. Myeloid cells and lymphocytes were further separated into different subtypes based on the same procedures. The singleR package (v1.4.0)(24) and “MouseRNAData” and “ImmGenData” reference databases were used to annotate the cell type of large cell populations. The cell types of clusters and subclusters were further confirmed and annotated by comparing the specifically expressed genes identified by the Seurat “FindAllMarkers” function with the known cell markers reported in the literature.

### Single-cell trajectory analysis and definition of cell states

The CD8^+^ T cell and memory likeT cell subpopulations of interest were selected for single-cell trajectory analysis. Using Monocle (v2.0)(25), cells were ordered according to their inferred pseudotime by following the steps described on Monocle documentation (http://cole-trapnell-lab.github.io/monocle-release/docs/). Only the top 100 differentially expressed genes identified by Seurat were used for dimensionality reduction and trajectory reconstruction. The reduce Dimension function with DDRTree as the reduction method was applied to the top principal components (PCs) and projected the cells onto two dimensions. After the dimension was reduced, the “orderCells” function was used to order cells and the plot_cell_trajectory function was used to visualize the trajectory in two-dimensional spaces.

### TCR sequencing data analysis

The clonotypes of single cells were defined using the Cell Ranger pipeline and default settings. TCR reads were aligned to mm10 reference genome. The consensus TCR annotation was performed using Cell Ranger VDR. Only in-frame rearrangement of TCR alpha and beta chains were considered to be productive and were used to define the dominant TCR of a single cell. Each unique dominant alpha-beta pair was defined as a clonotype. Two cells with identical alpha and beta sequences were assigned to the same clonotype. Cells harboring the clonotype that also presents in other cells were considered as clonal populations, and the number of such cells with the dominant alpha-beta pair indicated the degree of clonality of the clonotype. A total of 699 cells (young mice) and 356 cells (old mice) with such TCR alpha-beta pairs in old mice were identified.

### GO, KEGG, GSEA and GSVA analysis

GO and KEGG enrichment analyses were performed by WebGestalt(http://www.webgestalt.org), using genes specifically expressed in TEM-like cells as the input gene list. Gene Set Variation Analysis (GSVA) was used to estimate the enrichment scores of gene sets using the gene count data of each cell and was performed using R package “GSVA” (v1.37)(26). The differential pathways (log2 fold change > 0.32 and adjusted pvalue<0.05) identified by limma package (v3.44) were plotted in the bubble chart. GSEA analysis was performed for each cell subpopulation using the scaled gene expression matrix and GSEA package (v4.1) available at https://www.gsea-msigdb.org/gsea/downloads.jsp with default parameters. The gene sets for both GSVA and GSEA are provided by R package msigdbr.

### FACS Analysis

Single cells were obtained from draining lymph nodes and tumors of indicated mice. For cell surface analysis, a total of 1∼5 × 10^6^ cells were stained with Abs in the dark at 4°C for 30 min. After washing with cold FACS buffer (1 × PBS supplemented with 2% FBS), cells were analyzed using CytoFLEX flow cytometer (BECKMAN COULTER). CytExpert software was used for data analysis. To detect the expression of intracellular transcriptional factors, cells were fixed and permeabilized following 30-minute surface staining according to the manual of Foxp3 kit, followed by anti-Foxp3 antibody staining and FACS analysis. For cytokine analysis, cell samples were stimulated in vitro with PMA/Ionomycin in the presence of Brefeldin A (BioLegend) and Monensin (BioLegend) for 4 hours. Cells were washed and stained with surface marker antibodies, fixed and permeabilized using Fixation/Permeabilization buffer (BioLegend), and stained with intracellular antibodies.

### Quantification and statistical analysis

Statistical analyses were performed using GraphPad Prism (version 8) and R (version 3.6). Graphs were generated using GraphPad Prism and R ggplot2 package. Statistical analysis was applied to biologically independent mice or technical replicates for each experiment. The two-tailed Student’s t-test was used for all statistical calculations using GraphPad Prism 8 software. All error bars were reported as mean ± SEM with n=3 independent biological replicates. The level of significance is indicated as *P < 0.05, **P < 0.01, ***P < 0.001, ****P < 0.0001.

## Results

### Changes in immune cell composition of the TME during aging

To assess the effects of young and old host immune environment on tumor growth, we established syngeneic tumor models by subcutaneously implanting B16F10 melanoma cells into the right flanks of young (8-10weeks, n=3) and old (20-22months, n=3) C57BL/6J mice. Consistent with published results(5,12,27), old mice underwent significantly delayed tumor growth compared to young mice (**Fig. 1A**), indicating that host-intrinsic factors influenced tumor growth. To reveal the tumor-infiltrating immune cell composition, we profiled single-cell transcriptomes of CD45^+^ immune cells paired with TCR sequences of T cells in tumors isolated from young and old mice (**Fig. 1B**). After quality control and filtering, we obtained single-cell transcriptomes from 9,981 cells (4,606 for young mice and 5,375 for old mice) with a median of 973 genes (**Methods**). The single-cell RNA-seq (scRNA) data was normalized together and no batch effects were detected by PCA analysis (**Fig. S1A**). The clustering result was neither predominately contributed by cell cycle phases (**Fig. S1B**) nor by the sample group (**Fig. S1C**). Clustering analysis revealed 25 cell populations in the tumors of young and old mice. All clusters, except for cluster 25 expressing a high level of *Ptprc*, were identified as immune cells and used for subsequent analyses (data not shown). Analysis of the top 20 differentially expressed genes across these clusters revealed 10 cell types with unique transcriptional features (**Fig. 1C, Supplemental data 1**), including macrophages (*Adgre1*, *Cd14* and *Fcgr3*), conventional dendritic cells (*Xcr1*, *Flt3* and *Ccr7*), plasmacytoid dendritic cells (*Siglech*, *Clec10a* and *Clec12a*), monocytes (*Ly6c2* and *Spn*), neutrophils (*Csf3r*, *S100a8* and *Cxcl3*), natural killer cells (*Gzma*, *Klra4* and *Nkg7*), CD3^+^CD4^-^CD8^-^ T cells (*Cd3d, Cd3e* and *Cd3g*), CD8^+^CD4^-^ T cells (*Cd3d, Cd3e* and *Cd8*), CD4^+^CD8^-^ T cells (*Cd3g, Cd4* and *Ctla4*), and B cells (*Cd79a, Cd79b* and *Cd19*) (**Fig. S1D**). We visualized these cell types in two-dimensional spaces using t-distributed stochastic neighbor embedding (tSNE) and confirmed their cell identities by examining the expression of classic marker genes curated from the literature (28–30) (**Fig. S1E**). The distribution of cell types captured by scRNA was comparable between young and old mice (**Fig. 1D**). However, the proportions of CD3^+^CD4^-^CD8^-^ T lymphocytes, B lymphocytes, pDC, and neutrophils were dramatically reduced, whereas the proportions of macrophages, cDCs, and CD4^+^CD8^-^ T lymphocytes were increased in old mice compared to young mice (**Fig. 1E**), highlighting the significant differences of tumor-infiltrating immune cells during aging.

**Figure 1.**
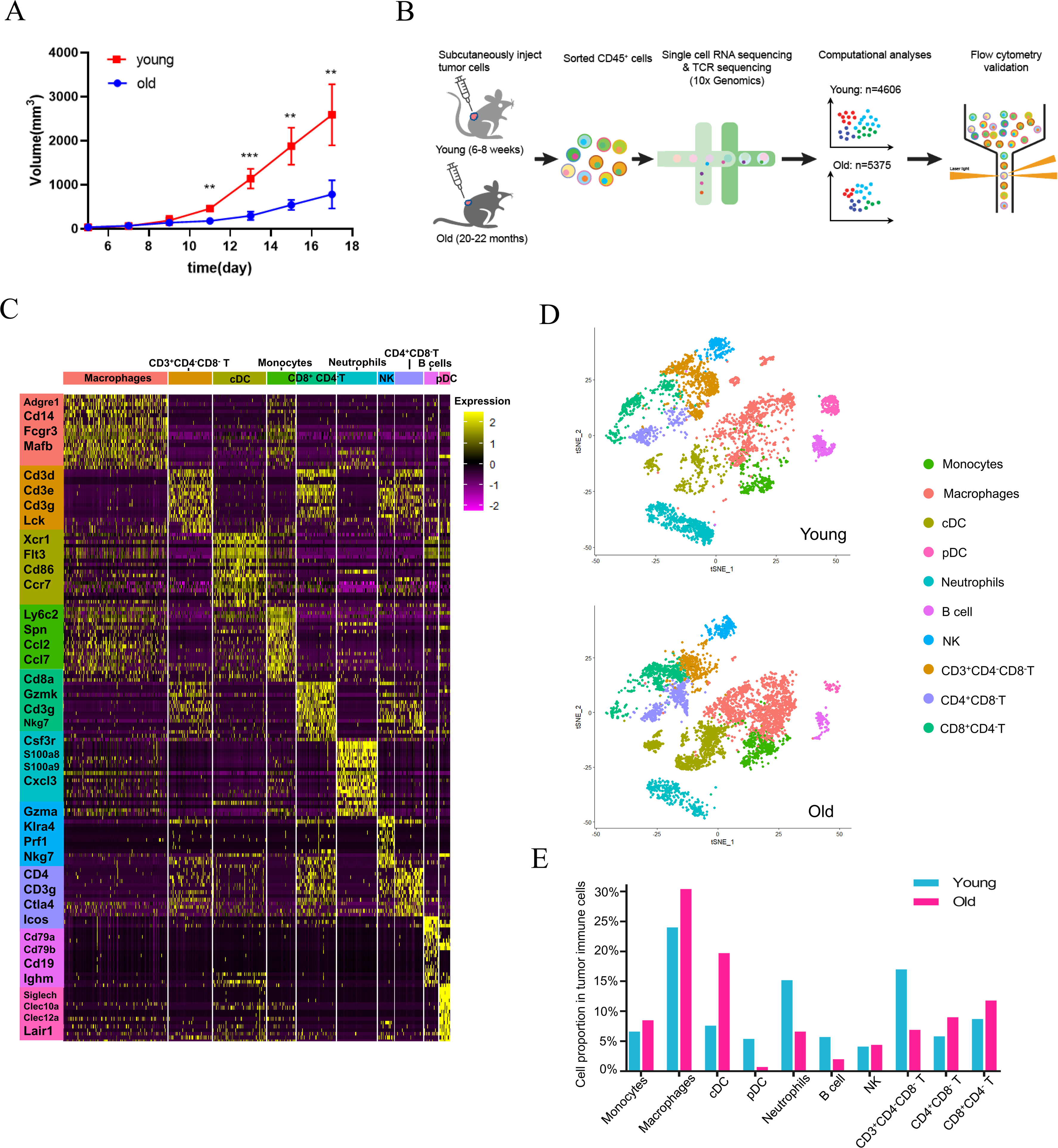
Changes in immune cell composition of the TME during aging (A) Tumor growth of B16 melanoma in young (n=3) and old (n=3) mouse models. Mice with the tumor size of (length × width^2^) / 2 mm^2^ were monitored. (B) Schematic diagram of the experimental design, single-cll RNA sequencing, data analysis, and validation. (C) Heatmap showing the relative expression (expression of normalized log2 (count +1)) of marker genes across different immune cell types. (D) tSNE projections of immune cells in tumors of young (upper) and old (below) mice. (E) The proportions of various immune cell types in tumors of young and old mice. cDC: conventional dendritic cells; pDC: plasmacytoid dendritic cells; NK: natural killing cells;

### Percentage comparison within tumor-infiltrating myeloid cells between young and old mice

Since myeloid cells were the most abundant cell type in our dataset, we first investigated the heterogeneity of myeloid cells in tumors of young and old mice through finer clustering. Although the size of total myeloid population was similar between young and old mice, old mice had significantly lower percentage of neutrophils but higher percentage of DCs in old mice in comparison to that of young mice. The proportions of tumor-infiltrating monocytes and macrophages remained unchanged (**Fig. 2A**). To assess if neutrophils had potential functional differences between young and old mice, we compared gene expression across two groups (**Fig. S2A**). Most genes displayed equivalent expression levels between young and old mice, suggesting that aging didn’t affect the functional activity of neutrophils which mainly relied on infiltration and cytokine production. Other myeloid cells, including macrophages, DCs, and monocytes, which consist of diverse subtypes (31, 32), were then partitioned into 5, 4, and 2 subsets, respectively (**Fig. 2B, S2B-C**). We confirmed their identities by examining the expression of classic markers (29, 33) (**S2D-E, Supplemental data 2**). The subsets that belong to the same cell type were clearly clustered together. Each subset was characterized by a specific gene expression pattern (**Fig. 2C**) and was labeled using their cluster index. Among them, the subtypes of macrophages and DCs showed striking differences in their proportions (**Fig. 2D**). We thus performed elaborate analyses on their cell subgroups next.

**Figure 2.**
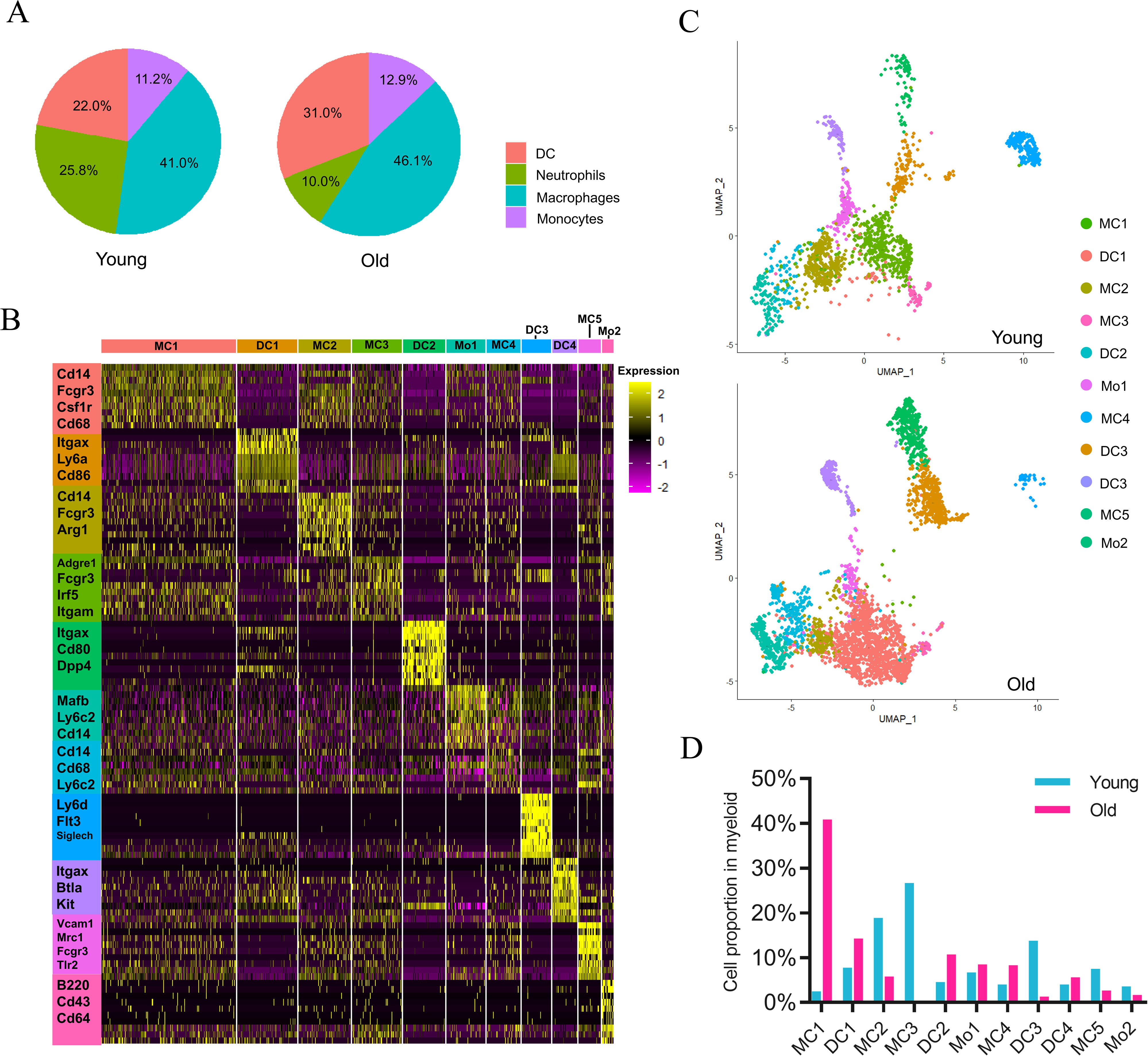
Myeloid cell composition of the TME in young and old mice (A) Pie charts showing the proportions of four major myeloid cell types in tumors of young and old mice. (B) Heatmap showing the relative expression (expression of normalized log2 (count +1)) of top differentially expressed genes across different myeloid cell clusters. (C) UMAP projections of myeloid cell subpopulations in tumors of young (upper) and old (below) mice. (D) The proportions of various myeloid cell clusters in young and old mice.

### Macrophage subsets in tumors of young and old mice

Macrophages are highly plastic cells that divide into diverse functional subsets in tumor enviroenment (31, 34). To understand the presence of macrophage subsets and their potential functions in the tumor immune microenvironment of old mice, we first evaluated the expression of classical M1-like and M2-like macrophage lineage markers in tumor-infiltriting macrophages (**Fig. 3A**). Most M1-like macrophage markers (e.g. inflammatory genes), including *Cd86*, *Cxcl10*, *Ccl9*, and *Ly6a*, were highly expressed in MC1, whereas the M2-like macrophage markers, i.e., *Mrc1* and *Arg1* were highly expressed in M2 and M4. All subsets showed considerable differences between two groups (**Fig. 3B**). The most striking observation was the accumulation of MC1 in tumors of old mice in contrast to highly enriched MC2 and MC3 in tumors of young mice.

**Figure 3.**
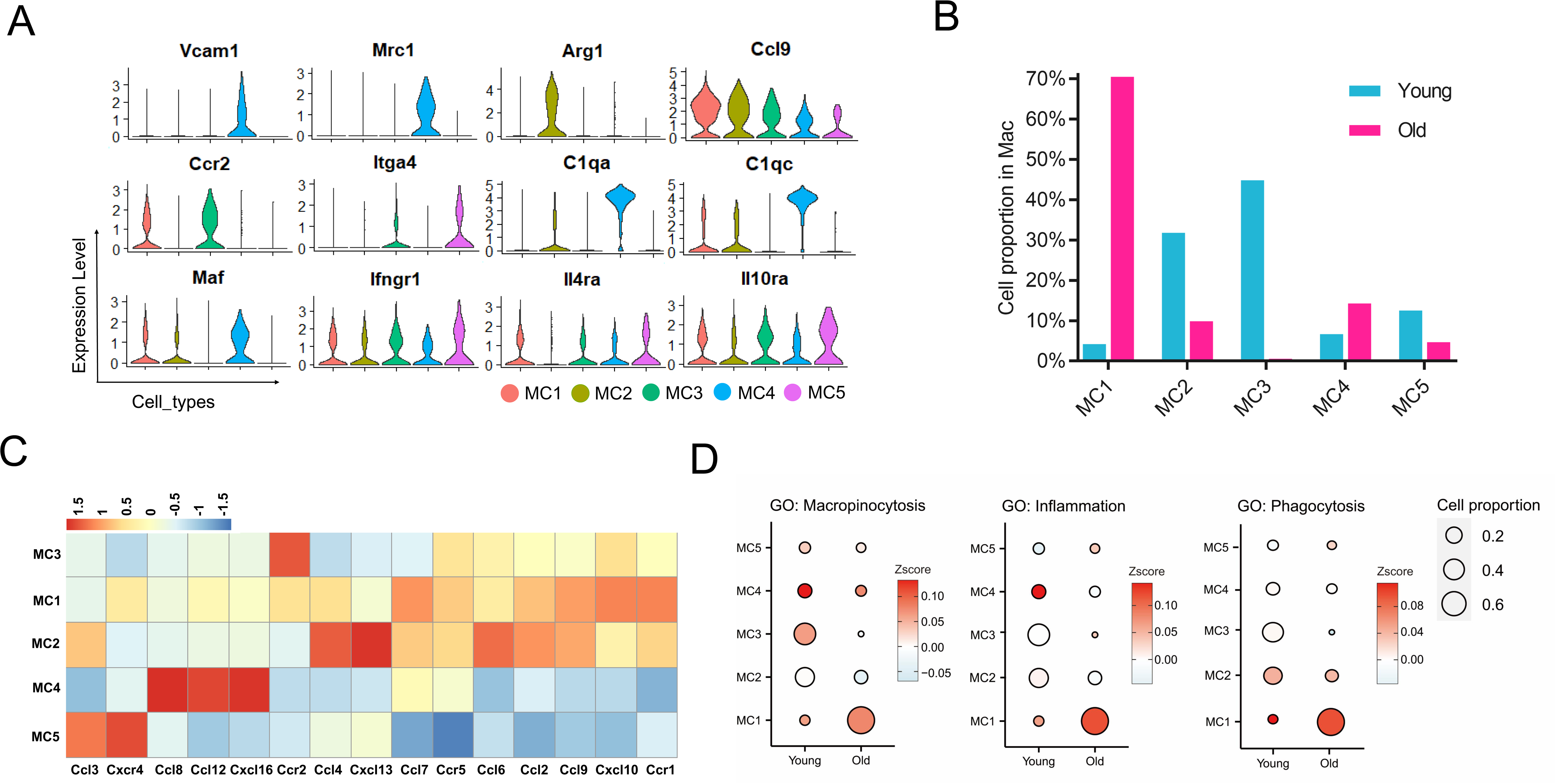
Macrophage cell subtypes and their heterogeneity in tumors of young and old mice (A) Heatmap showing the relative expression of marker genes across various macrophage subsets. (B) Comparison of the proportions of five macrophage subsets in young and old mice. (C) Expression profiles of chemokines in different macrophage subsets. (D) Bubble plots showing the scores (represented by the color gradient) of different gene sets and proportions (represented by the size of bubble) of each macrophage cluster in young and old mice. The gene set score is calculated by averaging the z-scores of gene expression values of all genes in this gene set. The gene expression in A & C is represented as expression of normalized log2 (count +1)). MC1 – MC5: five clusters of macrophages.

The chemokine system deeply affects activation, chemotaxis, and polarization of macrophages (35, 36). We found that each macrophage subset had a distinct chemokine expression profile (**Fig. 3C)**. MC1 expressed a high level of *Cxcl10* which may allow cells to attract both dendritic cells and T lymphocytes via chemotaxis (37, 38). MC2 highly expressed *Ccl4*, *Ccl6,* and *Cxcl13*; MC3 highly expressed *Ccr2*; MC4 highly expressed *Ccl8*, *Ccl12*, *Cxcl16,* and *Vcam1* which was associated with tumor progression (39); MC5 highly expressed *Ccl3*, *Cxcr4*, and integrin *Itga4* (also known as *Cd49d*), which are specifically expressed in the monocyte-derived TAMs (40). Moreover, we found that a large proportion of MC1 displayed the highest inflammatory, macropinocytotic and phagocytotic activities among all macrophage subtypes in old mice (**Fig. 3D, Supplemental data 3**). Interestingly, MC3 showed mild activity scores for most macrophage-associated pathways (**Fig. 3D).** Several genes including *Maf*, *Il4ra*, *Ly6c2*, and *C1qa* were significantly different in MC2 of young mice compared to old mice (**Fig.S3B**). Both MC2 and MC4 in old mice expressed higher Vegfa (**Fig.S3B, S3C**), which suggests that decreased angiogenesis in tumors of old mice was not due to decreased VEGFA production. Overall, we demonstrated that macrophages in old mice acquire increased pro-inflammatory states.

### Dendritic cell subtypes in tumors of young and old mice

Dendritic cells (DCs) are also heterogeneous and display various functional states in the TME (41). Four DC subsets were identified based on known biomarkers and genes critical for DC function (**Fig. 4A**). DC1 was identified as *Itgax*^+^*Itgam*^+^*Cd14*^+^ Monocyte-derived dendritic cells (MoDC); DC2 was identified as *Flt3*^+^*Zbtb46*^+^*Flt3*^+^*Ncoa7*^+^ conventional DC2 (cDC2); DC3 was defined as pDC with *Siglech*^+^*Irf8*^+^*Clec12a*^+^; DC4, which shared several common biomarkers with DC2 but specifically expressed *Clec9a*, *Btla*, *Xcr1*, and Itgae, was identified as cDC1. cDC2 and MoDC were highly enriched in tumors of old mice, while the pDC was highly enriched in tumors of young mice (**Fig. 4B**). We then determined the differential expression of genes relevant to cytokines production and chemotaxis, as well as those critical for antigen processing and presentation, and immune regulators, which allow DCs to regulate immune responses (**Fig. 4C, S4A, and S4B**). We showed that chemokine receptor *CCR7*, a member of the G protein-coupled receptor family responsible for the recruitment of lymphocytes and mature DC to lymphoid tissues, was highly enriched in MoDC; Both cDC1 and cDC2 highly expressed co-stimulatory genes *Cd80*, *Cd86*, *Cd83* and/or *Ly6a*, indicating their roles in activating local immune response. pDC highly expressed *Cd37*, *Lag3*, *Sla2*, and *Lair1*, suggesting their inhibitory roles in downregulating local immune response (**Fig. 4C**). We further confirmed the changes in pDC proportions by verifying the expression of *Lag3* in two groups using an independent cohort in both dLNs (**Fig. 4D**) and tumor tissues **(Fig. 4E)** of young and old mice. In addition, we found that high expression of the *H2* genes in cDC1 and MoDC suggests enhanced antigen presentation capacity (**Fig. 4SC**). To further explore the differences in subset proportions and potential functions between young and old groups, we performed gene GO analyisis on gene sets related to DCs (**Fig. 4F, Supplemental data 3**). We found that while pDCs in young mice produced larger quantities of cytokines, cDC1 and MoDC in old mice also tended to secrete more cytokines. cDC2 in old mice significantly expressed a batch of genes enriched in processes critical for dendritic cell differentiation, chemotaxis, positively regulation of antigen processing and presentation, while downregulating genes associated with dendritic cell apoptotic process. MoDC in old mice presented relatively higher ability in cytokine production, antigen presentiation and survival compared to that in young mice. Both *Fcgr1* and *Tcf4* genes, which are associated with immunoglobulin production, had higher expression levels in MoDC of old mice. In contrast, Cd37, which is associated with negative regulation of myeloid dendritic cell activation(42), was highly expressed in MoDC of young mice (**Fig. 4G**). Overall, these findings demonstrated that cDCs and MoDCs are prodominent populations in old mice supporting antitumor responses.

**Figure 4.**
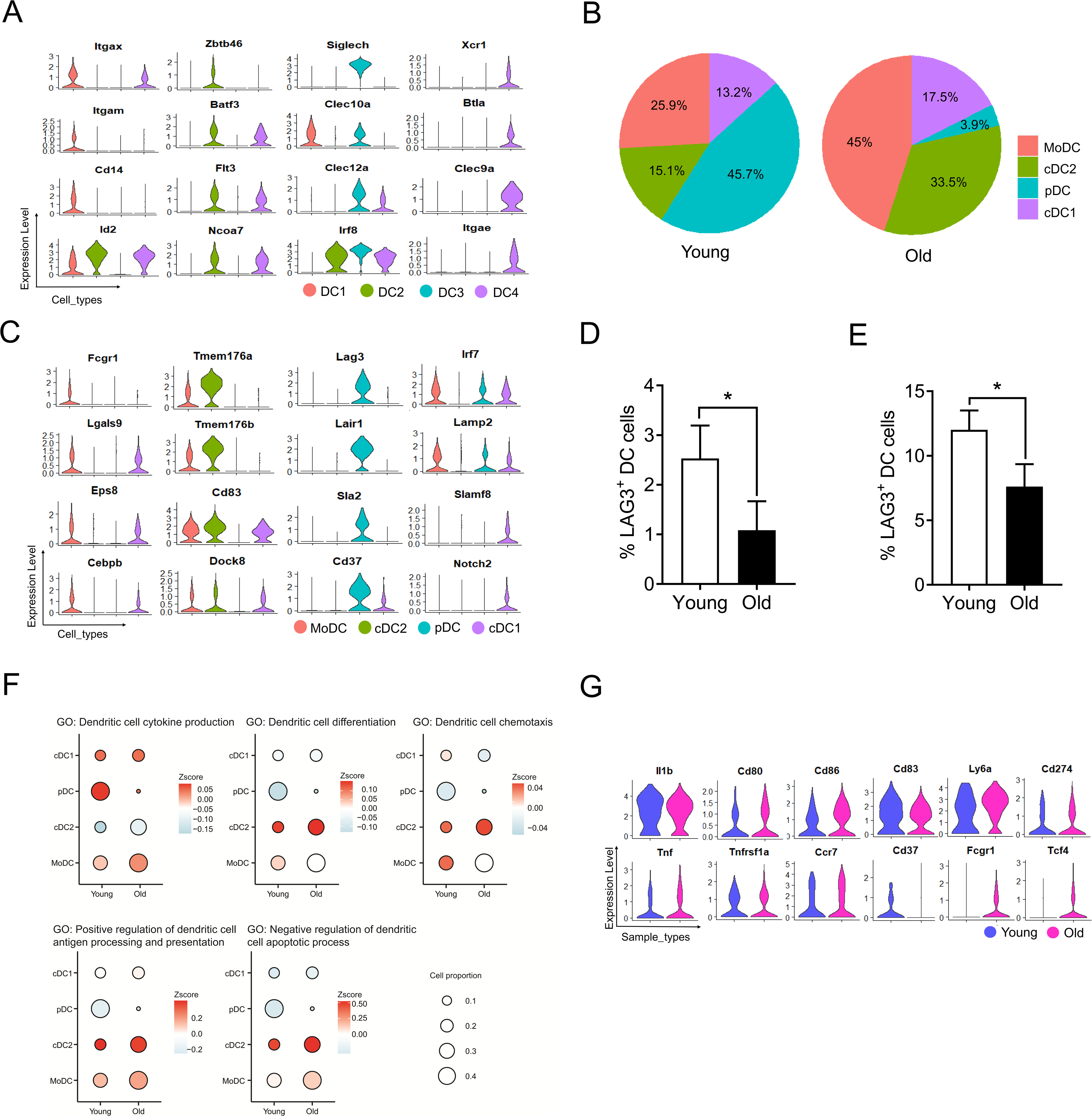
Dendritic cell subtypes and their heterogeneity in tumors of young and old mice (A) Violin plots comparing the expression levels of representative marker genes among different DC subtypes. (B) Pie charts showing the proportions of four DC subtypes in tumors of young and old mice. (C) Violin plots comparing the gene expression among different DC subtypes. (D and E) Percentages of Lag3^+^ DC in the dLN(D) and tumor tissues(E) of young and old groups(n = 3). (F) Bubble plots showing the scores (represented by the color gradient) of different gene sets and proportions (represented by the size of bubble) of each DC cluster in young and old mice. The gene set score was calculated by averaging the z-scores of gene expression values of all genes in this gene set. (G) Violin plot showing gene expression in MoDC in tumors of young and old mice The gene expression in A,C, and G is represented as expression of normalized log2 (count +1).

### Characterization of T cell subtypes in tumors of young and old mice

T cells are the main effector cells of adaptive immunity mediating antitumor response and their function is dependent on differentiation into distinct subsests (43). We extracted 2,934 T cells (1445 in young mice and 1489 in old mice) and performed unbiased clustering using highly variable genes to reveal 9 clusters of T cell populations. The clusters were visualized using uniform manifold approximation and projection (*UMAP*; **Fig 5A**). Each subpopulation was annotated based on the expression levels of the classical T cell markers (Fig**. 5B, S5A)** and regulators related to cell differentiation and function **(Fig 5C, S5B**). The universal T cell markers, e.g. *Cd3e*, *Cd3d*, were highly expressed in all subpopulations. Four clusters highly expressing *Cd8a* gene were assigned as CD8^+^ T cells, including activated_ CD8^+^ T (*Cd69*^+^*Cd28*^+^), exhausted_CD8^+^ T (*Gzmb*^+^*Prf1*^+^*Pdcd1*^+^*Lag3*^+^*Tigit*^+^*Havcr2*^+^), cytotoxic_CD8^+^T (*Gzmk*^+^*Gzmb*^+^*Pdcd1*^+^), and *Clec4e*_ CD8^+^ T (*Clec4e*^+^); two clusters were assigned as effector memory like (EM_like) with high expression of genes such as *Cd44*, *Cd69*, including CD8^+^ EM_like T with expression of *Gzmk* and *Cd8a,* and CD4^+^ EM_like T with expression of *Gzmb* and *Cd4*; The rest of clusters were assigned as naive_T with high expression of naïve gene markers *Ccr7*, *Sell*, *Tcf7* and *Lef1*, memory-like T cells (Memory_like T; *Il7r*^+^*Icos*^+^*Gzmk*^-^*Gzmb*^-^) and regulatory T cells (Treg; *Cd4*^+^*Foxp3*^+^) (**Fig. S5C**). The density distributions reflected difference in population frequency (**Fig. 5A, Supplemental data 4**). Old mice exhibited higher proportions of cytotoxic_ CD8^+^ T, CD8^+^ EM_like T, CD4^+^ EM_like T, and memory_like T but lower proportions of activated_ CD8^+^ T, exhausted_ CD8^+^ T, and naive_T compared with young mice (**Fig. 5D**). No obvious differences were observed in the proportions of Treg and Clec4e_ CD8^+^T cells between the two groups. In general, Tregs can be divided into two types with distinct phenotype and function, including naturally occurring Tregs (nTregs) and induced Tregs (iTregs). GSEA showed more genes associated with iTregs within tumors of young mice compared to old mice (**Fig. S6A**). Consistently, FACS analysis of Tregs in tumors of old mice revealed higher proportion of *Helios*^+^ cells (**Fig. S6B**) and higher level of *Helios* expression compared to that of young mice (**Fig. S6C**). *Ccr7*, controling thymic Tregs recirculation, was highly expressed in old mice (**Fig. S6D**). TGF-β, essential for Tregs to exert suppressive function (44), was also highly expressed in Tregs of old mice. Flow cytometry confirmed that old mice had significantly higher percentages of TGF-β^+^ Tregs in both dLNs (**Fig. S6E**) and tumor tissues (**Fig. S6F**). Clearly, more CD8^+^ effectors and similar Tregs were present in the TME of old mice.

**Figure 5.**
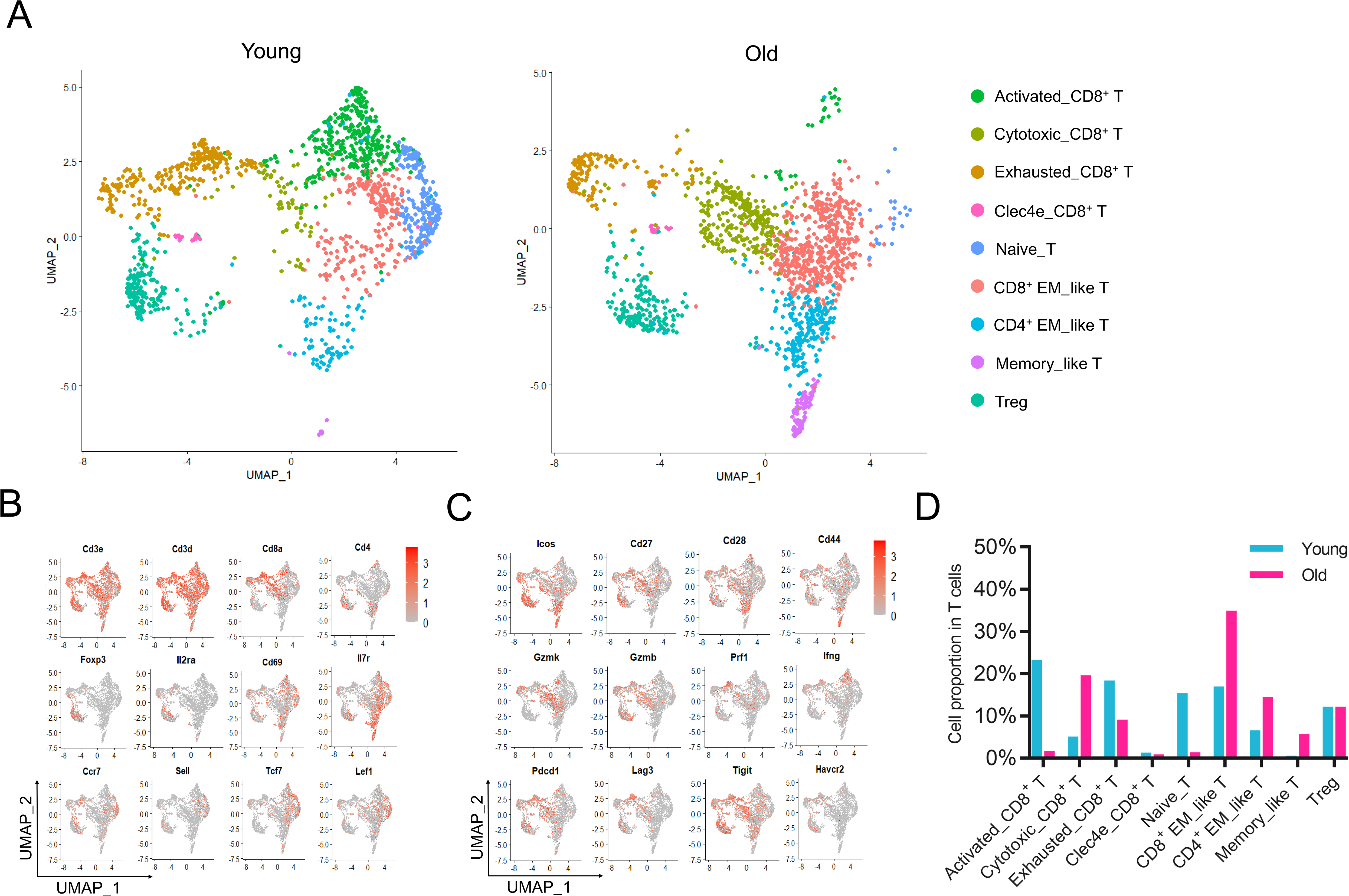
T cell subtypes and their heterogeneity in tumors of young and old mice (A) UMAP projections of T cells in tumors of young (left) and old (right) mice. (B and C) Expression of signature genes(B) and genes essential for T cell function(C) projected onto UMAP plots in(A). Color scale shows z-score transformation of log2 (count +1). (D) Bar graph showing the percentages of various T cell subtypes in the tumors of young and old mice.

### Striking shift of tumor-infiltrating CD8^+^ T subsets between young and old mice

To gain deeper insights into CD8^+^ T cell features in tumor environment of old mice, we further divided total CD8^+^ cells into four groups, including activated, cytotoxic, exhausted, and *Clec4e* positive CD8^+^ T cells. There was a higher proportion of cytotoxic CD8^+^ T cells in old compared to young mice while there were more activated CD8^+^ T cells and exhausted CD8^+^ T cells in young mice (**Fig. 6A**). Flow cytometry analyisis confirmed that old mice had lower percentages of PD-1^+^LAG3^+^ and Tim-3^+^LAG3^+^ (exhausted) cells (**Fig. 6B**), as well as higher percentages of *IFN*-γ ^+^ and Granzyme B^+^ (cytotoxic) cells in tumors (**Fig. 6C**). This result was also confirmed by the TCR repertoire analysis, where the top 10 clones in young mice were annotated as exhausted CD8^+^ T cells and the top 10 clones in old mice were annotated as cytotoxic CD8^+^ T cells (**Fig. 6D, Supplemental data 5**). To elucidate transcriptional fate of effector CD8^+^ T cells, we performed trajectory analysis on activated, cytotoxic, and exhausted CD8^+^ T cells (**Fig. 6E)**. The trajectory branch started with activated CD8^+^ cells and ended with cytotoxic and exhausted CD8^+^ cells (**Fig 6F, Fig S7B**), suggesting activated CD8^+^ T cells subsequently differentiated into cytotoxic or exhausted states. Moreover, we found that most genes associated with T cell exhaustion were enriched in young mice while the genes associated with T cell activation were enriched in old mice (**Fig. S7C**). Cytotoxic population were enriched in cell proliferation-related pathways in old mice whereas apoptosis process-associated pathways were enriched in young mice (**Fig. S7D**). The exhausted population expressed cell adhesion-related genes in old mice whereas pathways negative regulating proliferation were enriched in young mice (**Fig. S7E**). Consistent with pathway enrichment analysis, activated and exhausted CD8^+^ T cells in young mice displayed significantly higher apoptosis score (**Fig. S7F, Supplemental data 3**), suggesting effector T cells are more prone to death. Moreover, we analyzed shared TCR of exhausted population by all other CD8^+^ T cells, including activated CD8^+^ T, cytotoxic_ CD8^+^ T, and CD8^+^ EM_like T (**Fig. 6G**). The statistical results showed that tumor-infiltrated exhausted CD8^+^ T in young mice shared more TCR clones with activated_CD8^+^ T and cytotoxic_CD8^+^ T (**Fig. 6H**), suggesting that CD8^+^ T cells in young mice are more prone to exhaustion.

**Figure 6.**
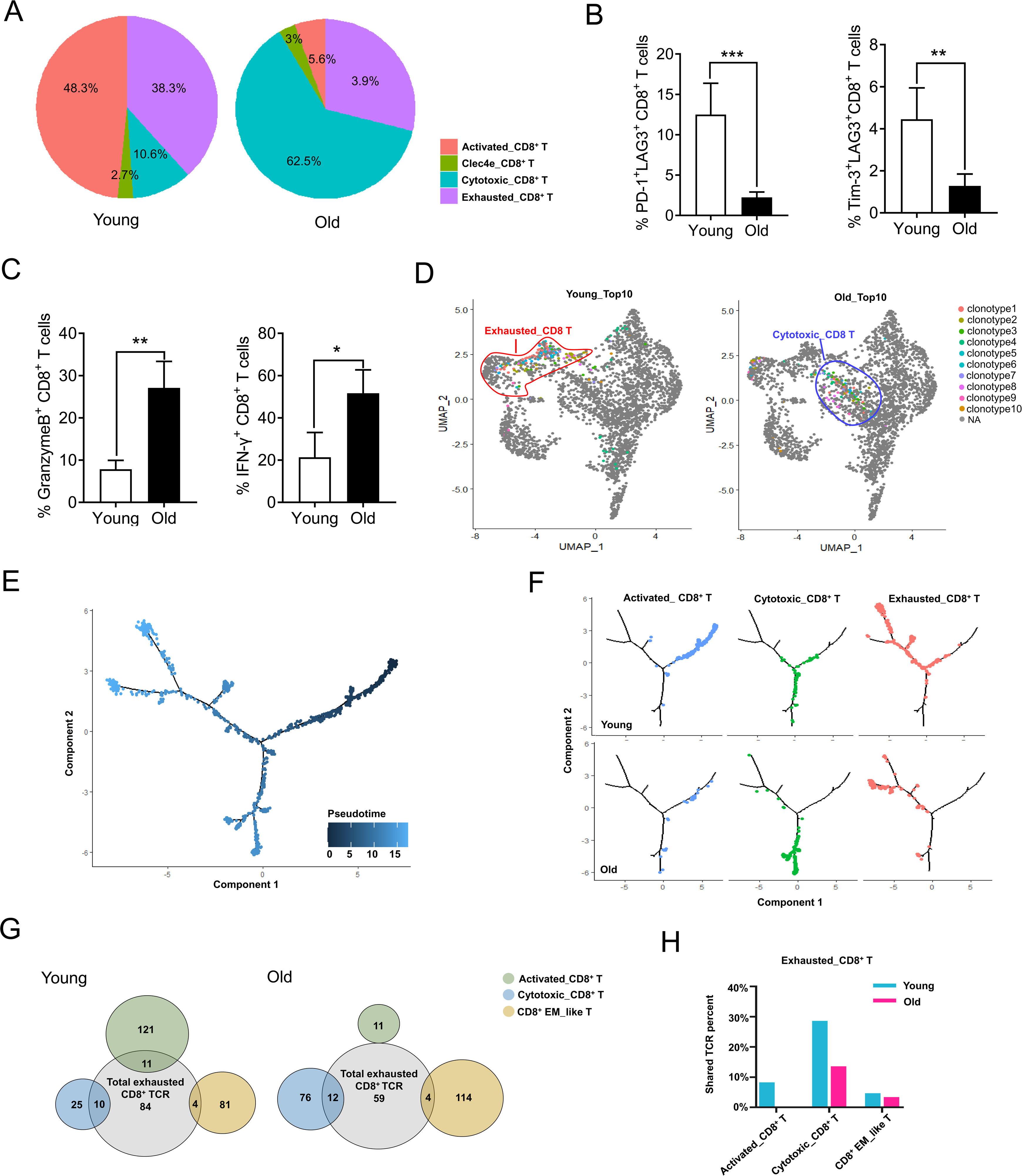
Diversity of tumor-infiltrating CD8^+^ T lymphocytes and their functional states in young and old mice (A) Pie charts showing the proportions of CD8^+^ T subtypes in young and old mice. (B) Percentages of PD-1^+^LAG3^+^ CD8^+^ T(left) and Tim-3^+^LAG3^+^ CD8^+^ T(right) cells in tumor tissue of young and old mice(n = 5). (C) Percentages of IFN-γ ^+^ CD8^+^ T(left) and GranzymeB^+^ CD8^+^ T(right) cells in tumor tissue of young and old mice(n = 3). (D) tSNE plots representing T-cell receptor (TCR) profiles of top10 clonotypes in tumors of young (left) and old (right) mice. (E and F) Differentiation trajectory of CD8^+^ T cells reconstructed by monocle2 using single-cell RNA-seq data. Color scale indicates either the ordering of cell in pseudotime (E) or the cell state (F). (G) Venn plot showing the number of shared TCR clones between exhausted CD8+ T and other CD8+ T cells in tumors of young and old mice. (H) Bar graph showing the percentages of TCR sequences shared by exhausted CD8+ T and other CD8+ T cells in young and old mice.

### Characterization of tumor-infiltrating CD8^+^ memory like T cells in young and old mice

In line with the previous reports that the number of memory T cells increase during aging in both human (45) and mice (46), we found that old mice exhibited higher proportions of CD8+ EM_like T CD4^+^ EM_like T, and memory_like T cells than young mice, as well as lower proportions of naive_T cells (**Fig. 7A**). Flow cytometry analysis confirmed more CD8^+^ TEM(**Fig. 7B**) and CD4^+^ TEM (**Fig. 7C**) cells in both dLNs(upper pannel) and tumor tissues(lower pannel) of old mice. While the expression of memory markers were similar, CD8^+^ EM_like T cells expressed higher levels of *Cd28*, *Ly6a* and *Gzmk* and lower levels of *Cd69*, *Fcer1g*, and *Cd7* in old mice compared to young mice (**Fig. S8A**). Similarly, CD4^+^ EM_like T cells expressed inflammation-related genes including *Nfkb1*, *Isg15*, and *Jund* in old group, whereas immunosuppressive genes including Tigit, and glycolysis-related gene *Aldoa* in young group (**Fig. S8B**).

**Figure 7.**
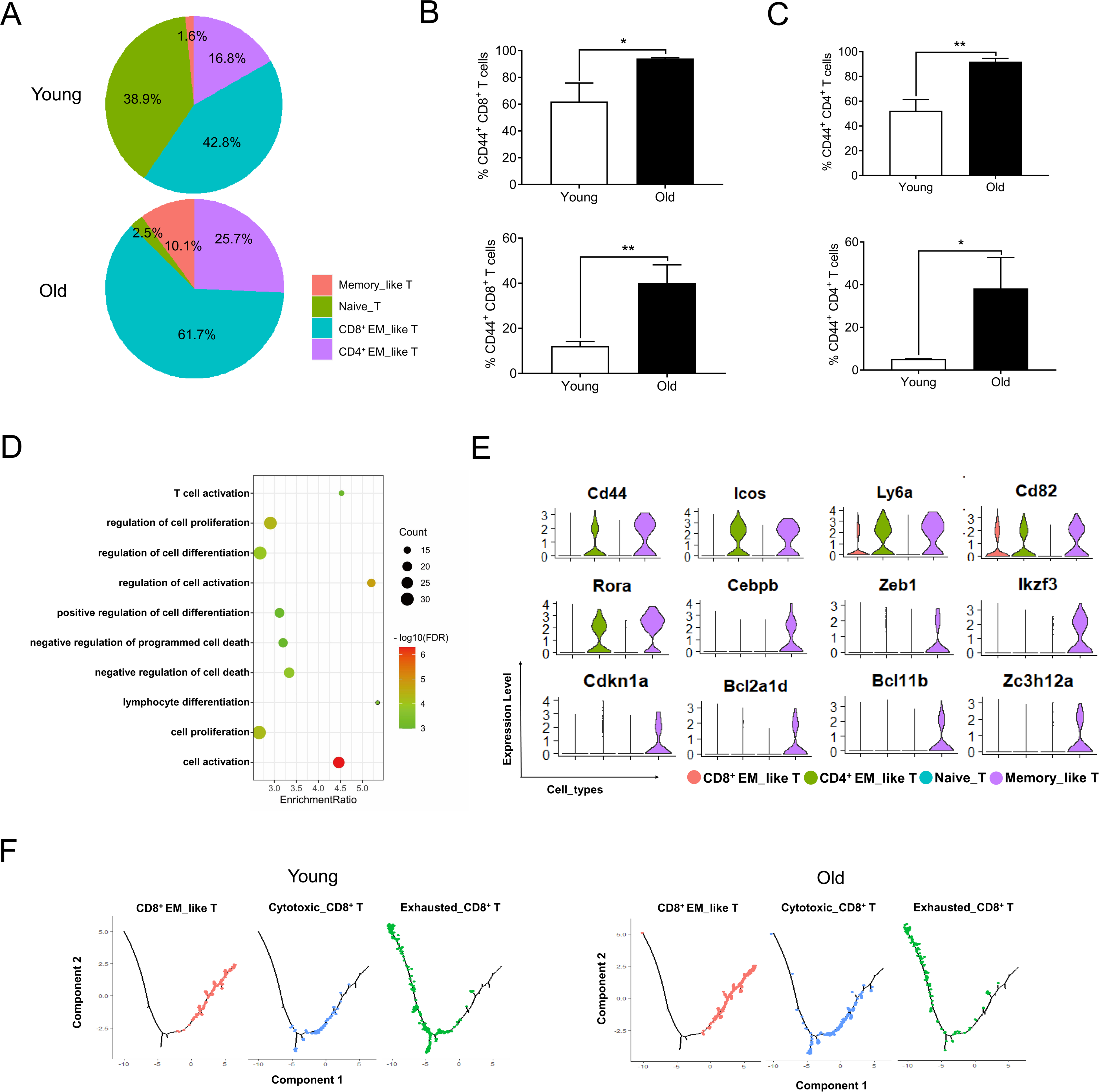
Characterization of tumor infiltrating naive and memory like CD8^+^ T cells in young and old mice (A) Pie charts showing the proportions of four T cells subtypes in young and old mice. (B and C) Percentages of CD8^+^ EM_like T(B) and CD4^+^ EM_like T (C) in the dLN (upper) and tumor tissues (below) of young and old. (D) Pathway enrichment result of top differential expressed genes in memory_like T cells. (E) Violin plot showing the gene expression in naive_T and different memory_like T cells. (F) Differentiation trajectory of CD8^+^ EM_like T cytotoxic_CD8^+^ T and exhausted_CD8^+^ T cells reconstructed by monocle2 using single-cell RNA-seq data from young and old mice. Colors indicate the cell differentiation states.

In addition to CD4^+^ and CD8^+^ EM_like T cells, we also discovered an undefined type of memory_like T cells, expressing genes related to T cell activation and differentiation (**Fig. 7D**). Apart from highly expressing costimulatory genes (*Cd82, Icos, Ly6a, Cd44*), memory_like T cells also expressed high level of differentiation-related genes (*Rora, Cebpb, Zeb1 and Ikzf3*), and proliferation-related genes (*Cdkn1a, Bcl2a1d, Bcl11b and Zc3h12a*) (**Fig. 7E**).

To further explain the differentiation relationship among CD8^+^ T cell populations, we constructed a trajectory tree for CD8^+^EM_like T, cytotoxic_ CD8^+^ T and exhausted_ CD8^+^T cells. CD8^+^ EM_like T cells were located at the beginning of the trajectory tree based on pseudotime (**Fig. S8C**), while cytotoxic_ CD8^+^ T, and exhausted_ CD8^+^T cells were located in the middle and ends of the trajectory tree, respectively (**Fig. S8D**). CD8^+^ EM_like T of old mice were activated to become cytotoxic_CD8^+^ T earlier than that of young mice. In contrast, cytotoxic_CD8^+^ T of young mice underwent exhaustion earlier and more readily than that of old mice **(Fig. 7F)**.

## Discussion

In this study, we reported the profile of immune cells in the TME of young and old hosts at the single-cell level and revealed possible immune mechanisms for old individuals that underwent delayed tumor progression. Given that most preclinical models used young animals, and most common cancers behaved differently in young and elderly patients, we and others (47–50) considered using old animals to explore the role of aging in the tumor immune microenvironment. Although the life span of mice and humans are very different, many studies have shown that humans and mice are very similar in disease progression, the physiological functions of various organs, and the molecular mechanism of aging (51, 52). In mice, senescence begins at least at 18 months. Therefore, we selected mice that are 20 to 22 month-old as the old group and 8 to 10 week-old mice as the young group. After confirming delayed tumor growth in old mice (47), we sorted CD45^+^ immune cells with flow cytometry before performing single-cell sequencing analysis. Our data showed that immune cells were distinct in composition and characteristics between these two groups, supporting that in comparison to young mice, old mice present a tumor environment with effector populations with stronger functional features to better defend against tumor progression.

T cell-mediated immunity is a vital component of the immune response for host defense against cancer (48). CD8^+^ T cells, the major cytotoxic killers of tumor cells, exhibited heterogeneity in the TME by single-cell sequencing (19,28,53,54). Here, we aimed at investigating tumor-infiltrating T cell populations and the corresponding characteristics, and determining the clonal dominance of CD8^+^ TILs in B16 melanoma tumor model. 9 types of T cells were characterized based on transcriptome analysis of single cells, including 4 types of CD8^+^ T cells, 3 types of memory like T cells, naïve T cells, and Tregs. We found that a large proportion of tumor-infiltrating CD8^+^ T cells were newly activated but exhausted in young mice, whereas a significantly higher proportion of cytotoxic or memory T cells are present in old mice. Flow cytometry further confirmed that old mice had higher percentages of IFN-γ ^+^ and Granzyme B^+^ CD8^+^ T cells in both dLNs and tumor tissues. Consistently, a recent study reported that Cd49d^hi^ Cd8^+^ cells are more abundant in tumors and exhibit a stronger anti-tumor effect in old mice (12). In addition, tumor-infiltrating CD8^+^ T cells showed enhanced survival in old mice, while those in young mice were more prone to apoptosis. The trajectory analysis of activated_ CD8^+^ T, cytotoxic_ CD8^+^ T, and exhausted_ CD8^+^ T cells showed that tumor-infiltrating CD8^+^ T cells underwent two sequential stages following activation with continuing differentiation trajectory in young mice, whereas CD8^+^ T cells were primarily located at the end of the trajectory tree in old mice, suggesting that effector CD8^+^ T cells underwent distinct differentiation programming during defense against tumor cells in the TME of young versus old mice.

With aging, continuous antigen stimulation and degeneration of the thymus cause the distribution of T cell subsets to shift from naive T cells to memory or memory-like cells (55). Previous studies demonstrated a reduction of response to neoantigens in elderly individuals due to a lower number of naive T cells (48). However, this does not necessarily indicate that the ability of old individuals to defend against foreign antigens is reduced since memory T cells cross-reactive to neoantigens may directly differentiate into effector T cells, and secrete inflammatory cytokines together with bystander memory-like cells. Consistently, there are several subsets of memory-like cells that exhibit stronger effector function. In old mice, CD8^+^ EM_like T expressed higher levels of costimulatory genes: *Ly6a*, *Cd28*, and the cytotoxic gene *Gzmk,* while CD4^+^ EM_like T expressed higher levels of inflammation-related genes: *Nfkb1*, *Isg15*, and *Jund*. In contrast, CD4^+^ EM_like T cells in young mice expressed higher inhibitory genes: *Tight*, and *Aldoa*. In addition, we also identified a unique population, Memory_like T cells, which highly expressed *Il7r*, *Cd44,* and co-stimulation genes (e.g. *Cd82*, *Icos*, *Ly6a*). Gene expression enrichment analysis showed that upregulated genes in these cells were related to cell differentiation, proliferation, and T cell activation, suggesting that this cluster of unique T cells in the elderly represent recently activated effector cells derived from memory T cells. Analysis of shared TCR clones between T cell populations in different states can be used to infer differentiation and cell origin(56). Compared with the old group, CD8^+^ EM_like T, cytotoxic_CD8^+^ T cells, and exhausted_CD8^+^ T cells in the young group shared a higher proportion of TCR clones. The relationships among 3 subpopulations were further confirmed by trajectory analysis.

NK cells are another key player of immune surveillance against tumorgenesis. However, different from the general decline of T cells with aging(57), the proportion and function of NK cells remained relatively stable. Although GSEA showed that NK cells of young mice were relatively active(**Fig. S9A**), many functions showed no difference except for pathways related to metabolism (**Fig. S9B**). Our single-cell data demonstrated that tumor-infiltrating NK cells expressed similar levels of functional molecules, such as *Prf1*, *Gzmb*, *Nkg7*, *NKG2D*(*Klrk1*), Ly-*49g*(*Klra7*), and *CD94*(*Klrd1*) in both young and old hosts (**Fig. S9C**). Consistent with a previous report in humans(58), our data also showed that NK cells in old mice expressed a lower level of *NKG2A*(*Klrc1*) but a higher level of *Ly49*(*Klra4*) (**Fig. S9C**). In addition, we found that NK cells in old mice expressed higher levels of inflammatory genes (*Nfkb1* and *Stat3*) and activation molecule 4-*1BB* (59), indicating stronger effector function of NK cells and the proinflammatory status in the elderly.

Previous studies show that B cells are generated at a reduced rate(60), showing lower levels of percentage and absolute numbers(61, 62) while also differentiating into longer-lived memory B cells(62) in old individuals. In the tumor environment, our data also showed that the infiltrating B cells in the tumors of old mice tended to be memory B cells (**Fig. S10A**), and were present at a lower percentage than that in young mice (**Fig. S10B**). In addition, tumor-infiltrating B cells of old mice expressed lower levels of canonical B cell markers (*Cd19*, *Cd79a*, and *Cd79b*) compared to that of young mice (**Fig. S10C**). These cells were similar to a type of exhausted or “double negative” memory B cells expressing low level of IgD and *Cd27* reported in a previous study (63). IgM memory B cells accumulate with age and become the predominant memory B cell subset. Consistently, our data showed that tumor-infiltrating B cells in old mice express higher level of IgM but lower IgD expression. Previous studies found that 4-*1BBL*-expressing B cells increase with age in humans and mice, enhancing Granzyme B expression of CD8^+^ T cells by presenting endogenous antigens(12), however, the current study did not reveal an enrichment of this population in the tumor of old mice.

Among myeloid cell lineages, cDCs and M1-type macrophages play positive roles in defending against tumor growth through innate mechanisms and activation of adaptive immunity (31, 32). cDC1 mainly activates CD8^+^ T cells by processing and presenting exogenous antigens through MHC class I molecules, while cDC2 is crucial for inducing the immune response of CD4^+^ T cells (32, 64). Our data also showed that the proportion of dendritic cells that produce the inflammatory factor (TNF-α), and *Ccl5*, critical for cDC1 recruitment to TME(65), were increased in old mice. cDC2, vital to stimulating Cd4^+^ T cell-mediated immunity in cancer (66), had similar abilities of antigen processing and presentation in young and old mice, but a higher population of cDC2 in old mice. M1-type macrophages express higher inflammatory genes and are associated with antigen presentation(34, 67). We showed that a higher percentage of macrophages in the tumor of old mice are M1-type with enhanced inflammatory factor expression (TNF-α) and potential costimulatory function (*Cd80*, *Cd86*). Previous studies showed that M1-type macrophages in old mice display powerful phagocytosis and micropinocytosis abilities, which are beneficial to the internalization of exogenous antigen(68).

In parallel to effector cells playing positive roles in mediating anti-tumor response, there are well-identified immune cells that negatively regulate immune response, including Tregs, MDSCs, M2-type macrophages, and pDCs in tumor environment(69, 70). Previous studies revealed that elderly individuals demonstrated changes in Tregs’ frequency and function with aging (71, 72). In contrast, the current study showed that the proportion and functional gene expression of Tregs in old mice were similar to that in Tregs of young mice. However, tumor-infiltrating Tregs tended to be of thymus origin (nTregs) in old mice whereas more are iTregs in young mice, supported by flow cytometry analysis showing higher expression of *Helios* in Tregs from old mice. In addition to Tregs, MDSCs are specialized immunosuppressors to prevent excessive inflammation (73, 74) and promote tumor growth and metastasis(33). However, no MDSCs were identified in the current data. M2-type macrophages are characterized by high expression of *Arg1, Vcam1, or Cd206*, and are associated with immunosuppression (75) and tumor growth (31). We identified two types of macrophages with high expression of *Arg1*(MC2) and high expression of *Vcam1*(MC4), and both groups of macrophages were elevated in young mice. pDCs have been shown to associated with poor patient prognosis in several cancers by inducing the expansion of Tregs(76). pDCs were present a lower level in old mice, potentially allowing for more robust anti-tumor immunity.

In conclusion, we utilized single-cell RNA sequencing analysis to evaluate the proportion and gene expression difference of immune cell components in the tumors of young and old mice. In comparison to young mice, old mice showed a higher proportion of effector cells (cytotoxic T cells, cDC, and M1-type macrophages) and a lower proportion of dysfunctional and immunosuppressor cells (exhausted T cells, iTregs, pDC, and M2-type macrophages) in the TME. TCR spectrum and trajectory inference analysis demonstrated preferential differentiation of CD8^+^ T cells into cytotoxic cells in old mice versus exhausted cells in young mice. However, the underlying mechanisms that regulate distinct differentiation paths requires further investigation.

## Supporting information

The top differentially expressed genes of each cluster for all immune cells

The top differentially expressed genes of each cluster for myeloid cells

The different cell gene signatures

The top differentially expressed genes of each cluster for T cells

Top10 TCR clones

## Code availability

The codes generated in this study are available at the Github repository (https://github.com/Immugent/single-cell-data-process)

## Data Availability Statement

The raw data will be made available by the authors, without undue reservation, to any qualified researcher.

## Ethics Statement

The animal study was reviewed and approved by the Animal Care Committee of Xi’an Jiaotong University.

## Author Contributions

B.Z. and C.Z. conceptualized the study and designed experiments. C.Z. and H.Z. collected the tumor samples with help from L.L., Y.S., H.L, and X.W. L.S., A.J., Y.Z. and C.Z. prepared samples for single-cell sequencing and performed experiments with help from X.Y. and Y.S. C.Z. analyzed scRNA-seq data with input from Z.X. C.Z. and B.Z. wrote the manuscript. B.Z., Z.X., C.S., X.Z. and Y.H. reviewed and revised the manuscript.

## Acknowledgments

We thank Dr. Guohua Zhang from Beckman Coulter, Prof. Chen Huang and Dr. Xiaofei Wang from Department of Cell Biology and Genetics for flow cytometric technical supports.

## Supplementary Figure legend

**Fig. S1.**
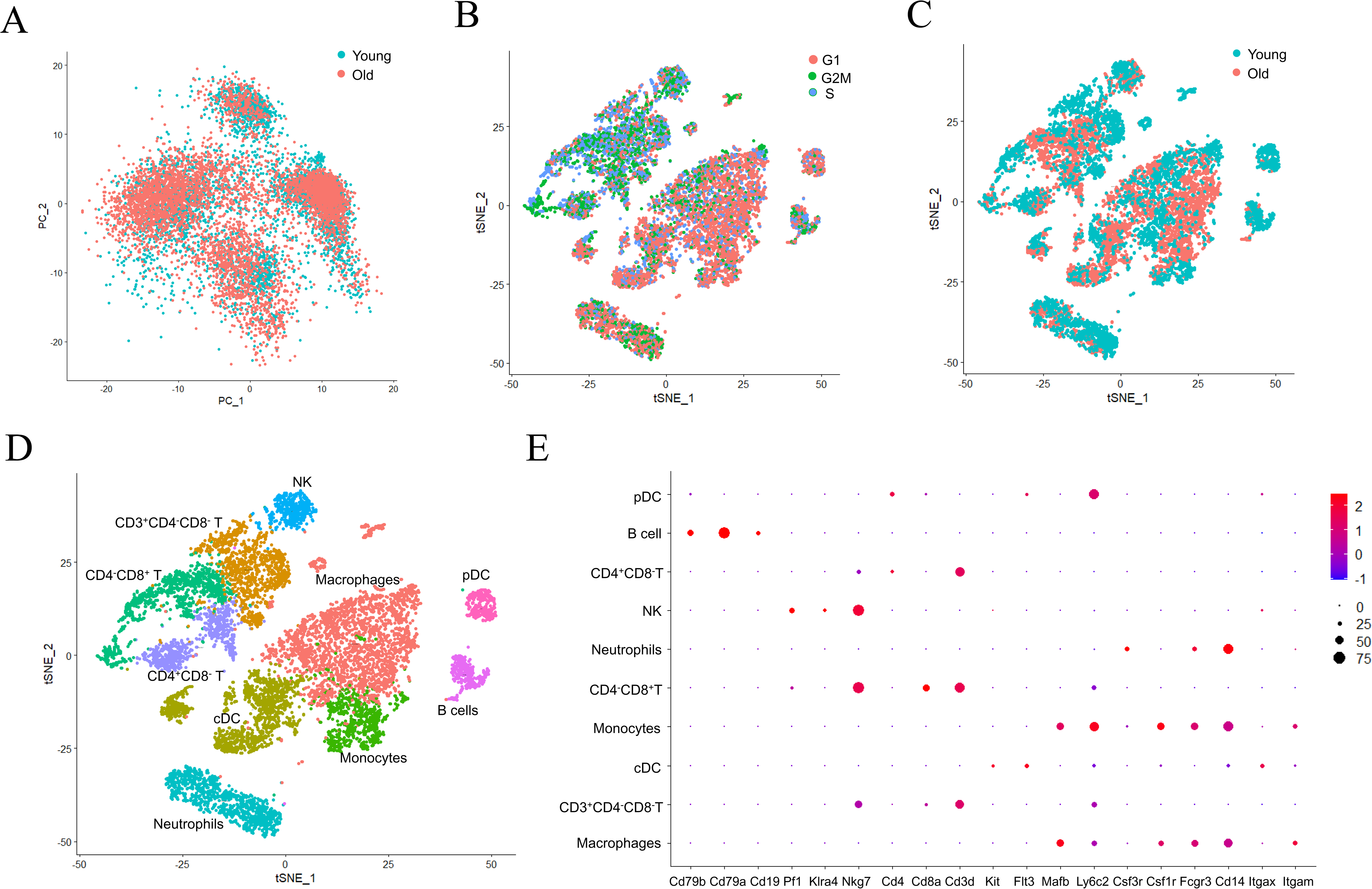
scRNA-seq profile of tumor-infiltrating immune cells in young and old mice (A) PCA plot of 9981 high-quality immune cells showed no batch effect existing between young mice and old mice. (B) tSNE visualization of the cell-cycle scores for different immune cells (C) tSNE plot of the distribution of all immune cells, colored by age groups. (D) tSNE plot of all immune cells from tumors of both young and old mice. (E) Dot plot showing the expression levels of marker genes in different cell clusters. The expression was measured as the log_2_ (count+ 1).

**Fig. S2.**
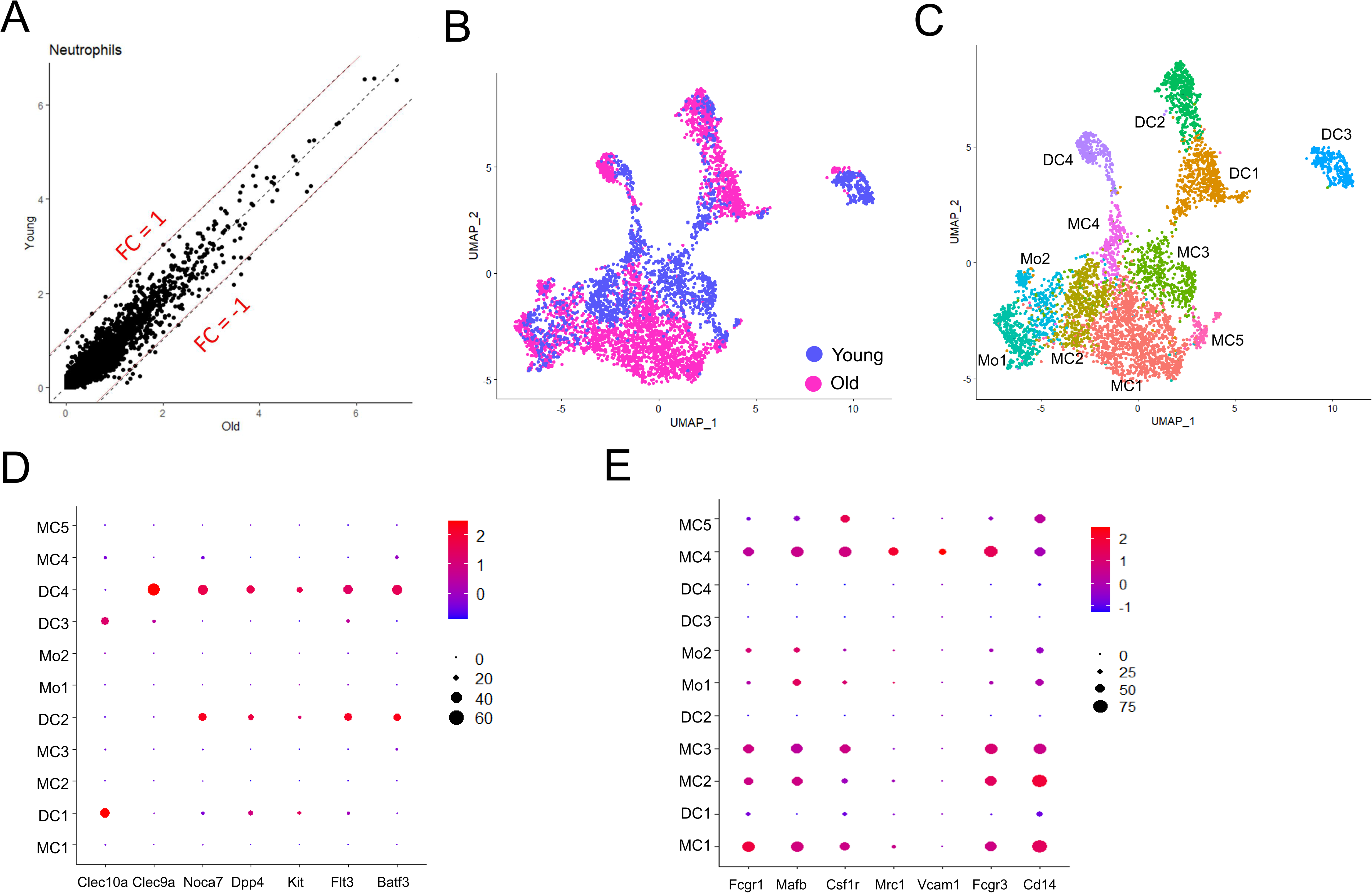
Characterization of myeloid cells using scRNA-seq data (A) Scatter plot comparing the gene expression profiles of neutrophils in tumors of young and old mice. Each dot denoted an individual gene. The criteria for gene changes are |FC| >1. (B) UMAP plot of macrophages, dendritic cells, and monocytes, colored by age groups. (C) UMAP plot of macrophages, dendritic cells, and monocytes in tumors of both young and old mice. (D and E) Dot plot showing the expression levels of representative DC markers(D) and macrophage markers(E) of in different cell clusters. The expression was measured as the log_2_ (count + 1).

**Fig. S3.**
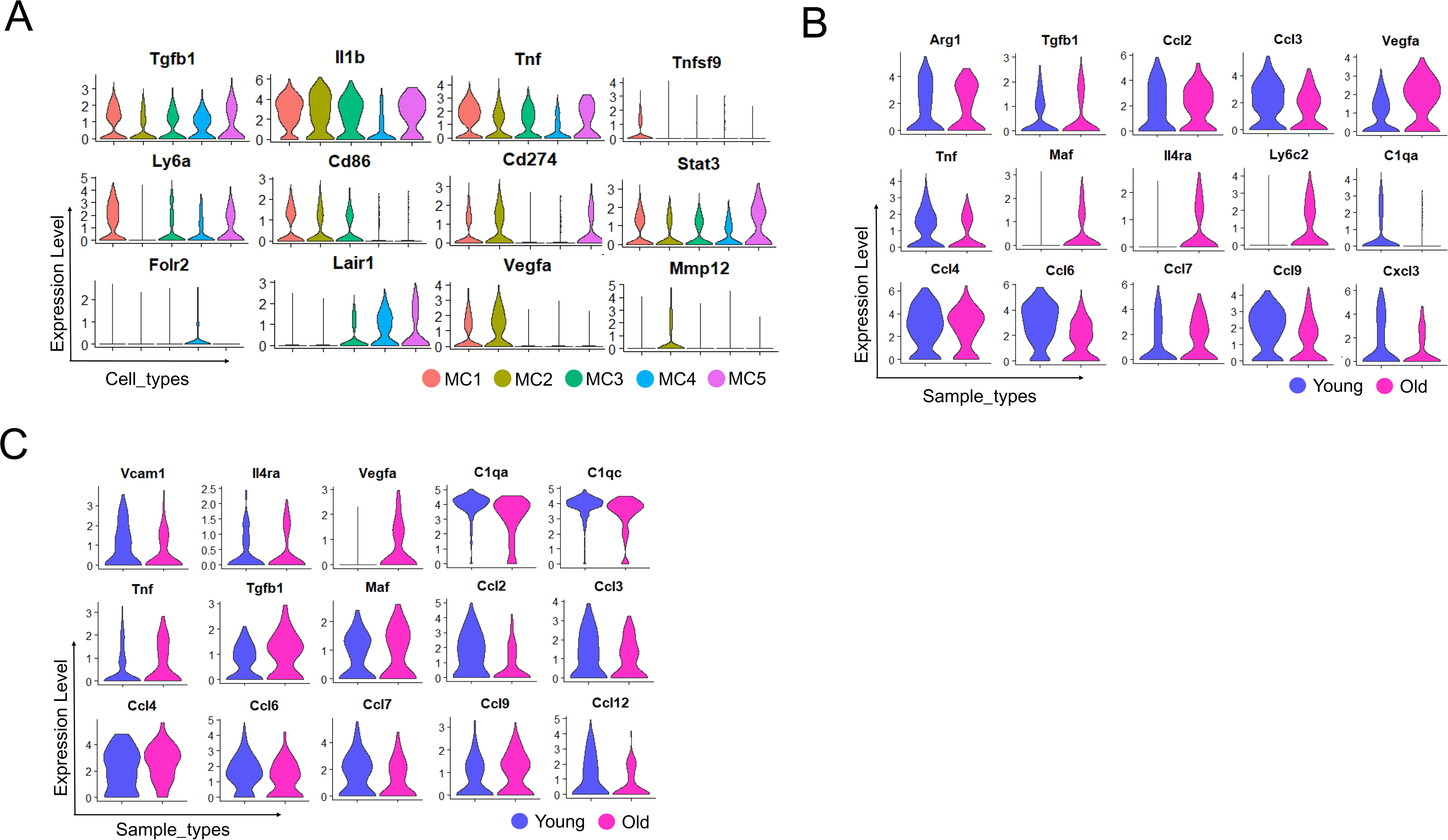
Characterization of tumor infiltrating macrophages Violin plot comparing the expression levels of representative signatures among different macrophage subtypes. (B and C) Violin plot comparing the expression levels of representative MC2 signatures(B) and MC4 signatures(C) in young and old mice. The gene expression was measured as the log_2_ (count + 1). markers(E) of in different cell clusters. The gene expression was measured as the log_2_ (count + 1).

**Fig. S4.**
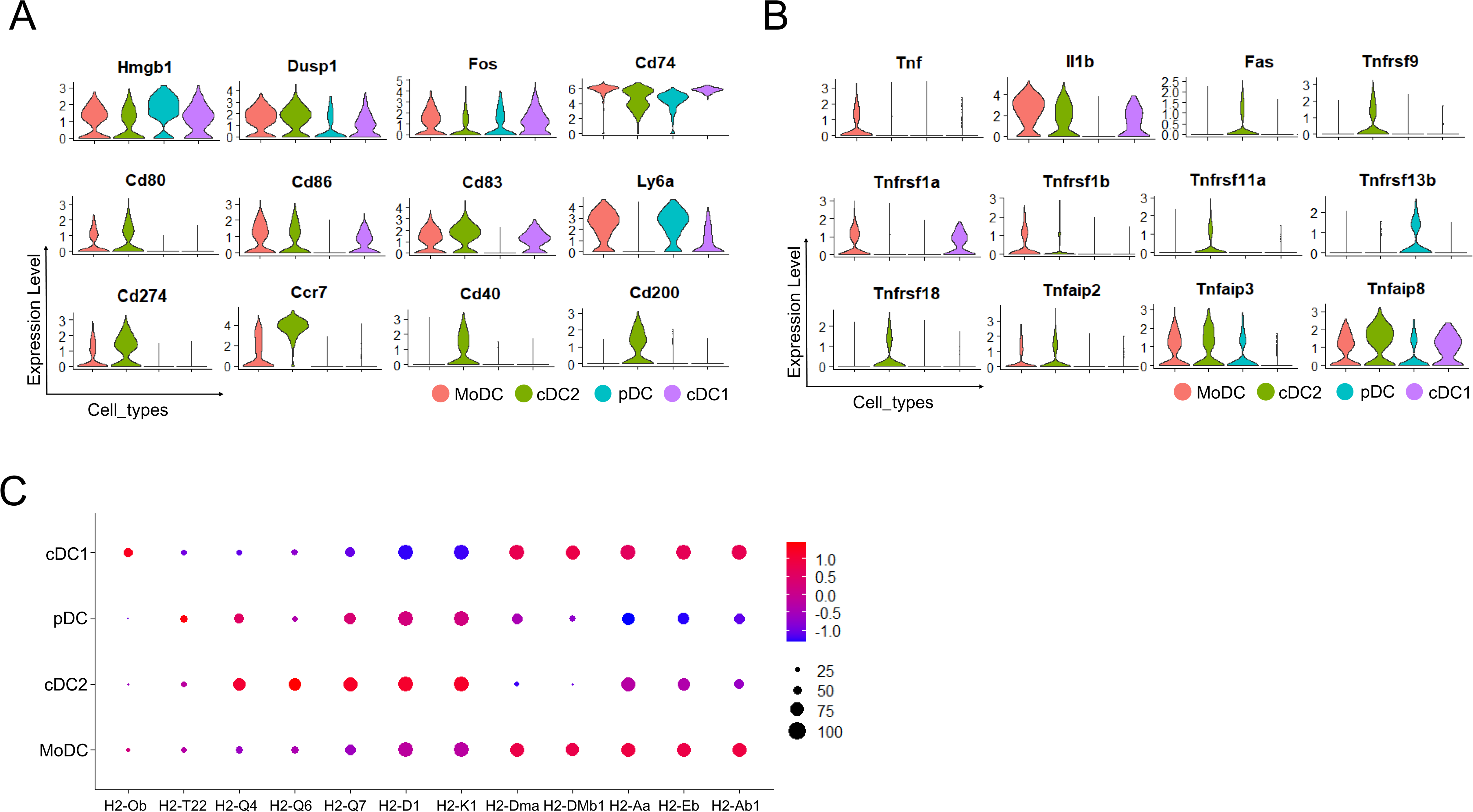
Characterization of tumor infiltrating dendritic cells (A) Violin plot comparing the expression levels of functional genes among different dendritic cell subtypes. (B) Violin plot showing the expression of Tnf-family genes in various dendritic cells. (C) Dot plot showing the gene expression of H2 components in various dendritic cells. The gene expression was measured as the log_2_ (count + 1).

**Fig. S5.**
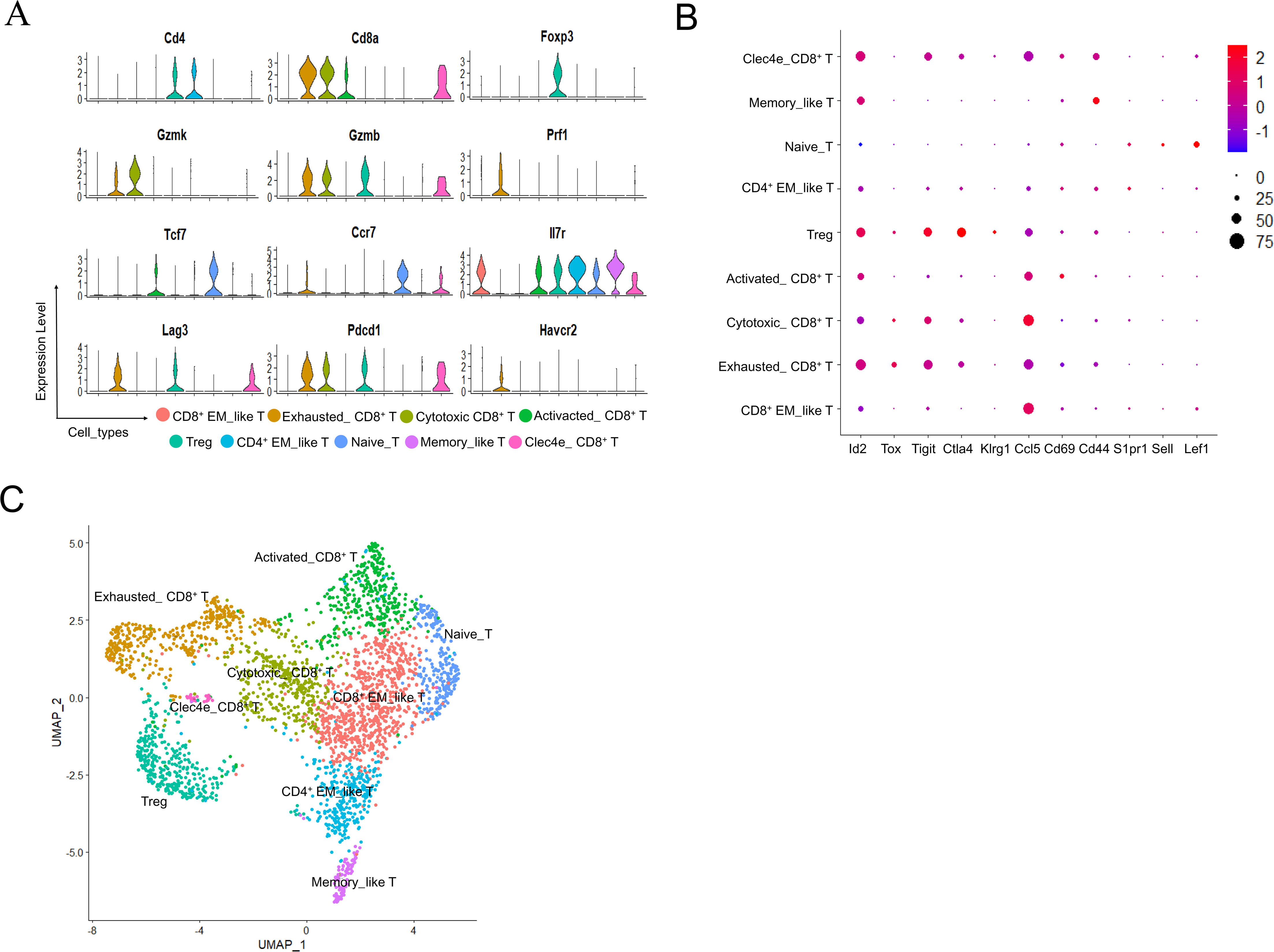
Characterization of tumor infiltrating T cells (A) Violin plot showing expression levels of representative marker genes for different T cells. (B) Dot plot showing the expression levels of representative marker genes of different T cells. (C) UMAP plot showing the distribution of different tumor infiltrating T cells, colored by the cell subtypes. The gene expression was measured as the log_2_ (count + 1).

**Fig. S6.**
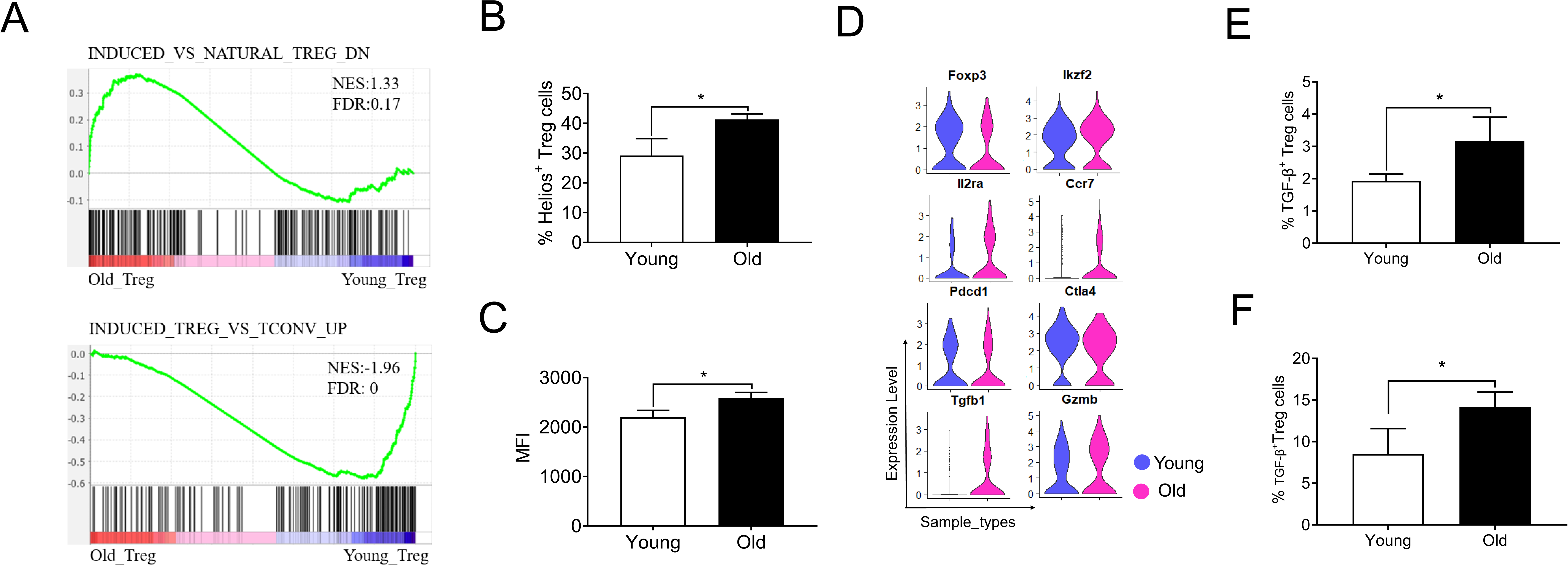
Analysis of tumor infiltrating Tregs (A) GSEA plots of the signature gene sets in tumor infiltrating Tregs of young mice vs. old mice. NES: Normalized Enrichment Score. (B) Percentages of tumor-infiltrating nTregs (Helios^+^) in the young and old mice (n = 3). (C) Histogram of Helios expression in young and old Tregs(n = 3). (D) Violin plot showing expression levels of Tregs functional genes in young and old mice. The expression was measured as the log_2_ (count + 1). (E and F) Percentages of TGF-β^+^ Tregs in the dLN(E) and tumor tissues(F) of young and old mice(n = 3). Error bar is mean ± SEM. The significant level is determined by two-tailed, paired Student’s t-test.

**Fig. S7.**
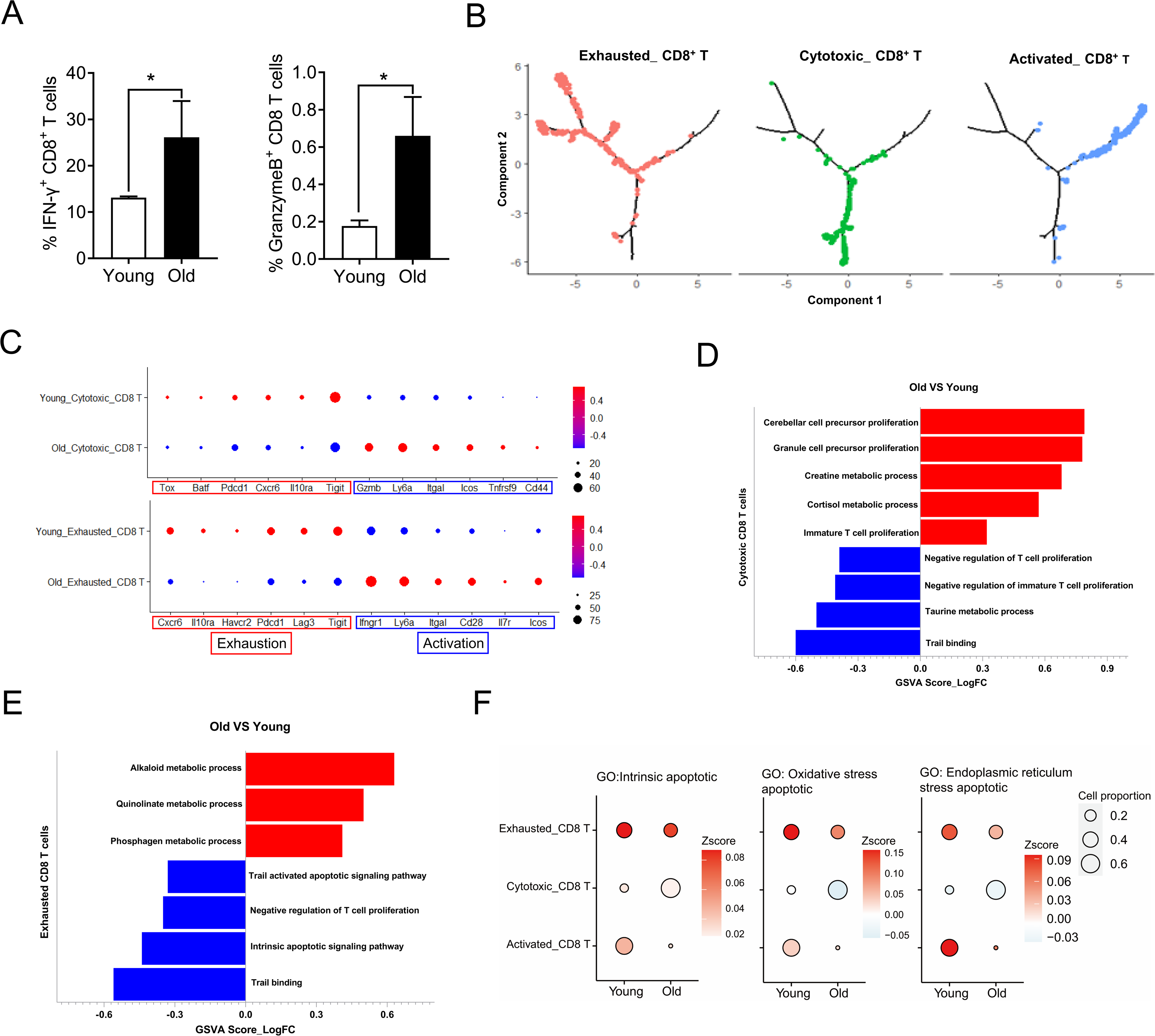
Analysis of CD8^+^ T cell differentiation status (A) Percentages of IFN-γ CD8 T(left) and GranzymeB CD8 T(right) cells in dLN of young and old mice(n = 3). (B) Cell differentiation trajectory of activated CD8^+^ T cells (right), cytotoxic CD8^+^ T cells(middle), and exhausted CD8^+^ T cells (left) reconstructed by Monocle2 using single-cell RNA-seq data. Color scale indicates the ordering of cells in pseudotime. (C) Dot plot showing expression of genes associated with T cell states in cytotoxic CD8^+^ T cells(up) and exhausted CD8+ T cells (down). (D and E) Gene Set Variation Analysis (GSVA) results using differential expressed genes in cytotoxic CD8^+^ T cells(D) and exhausted CD8^+^ T cells(E). (F) Bubble plots showing the scores (represented by the color gradient) of different gene sets and proportions (represented by the size of bubble) of each T cell groups in young and old mice. The gene set score was calculated by averaging the z-scores of gene expression values of all genes in this gene set.

**Fig. S8.**
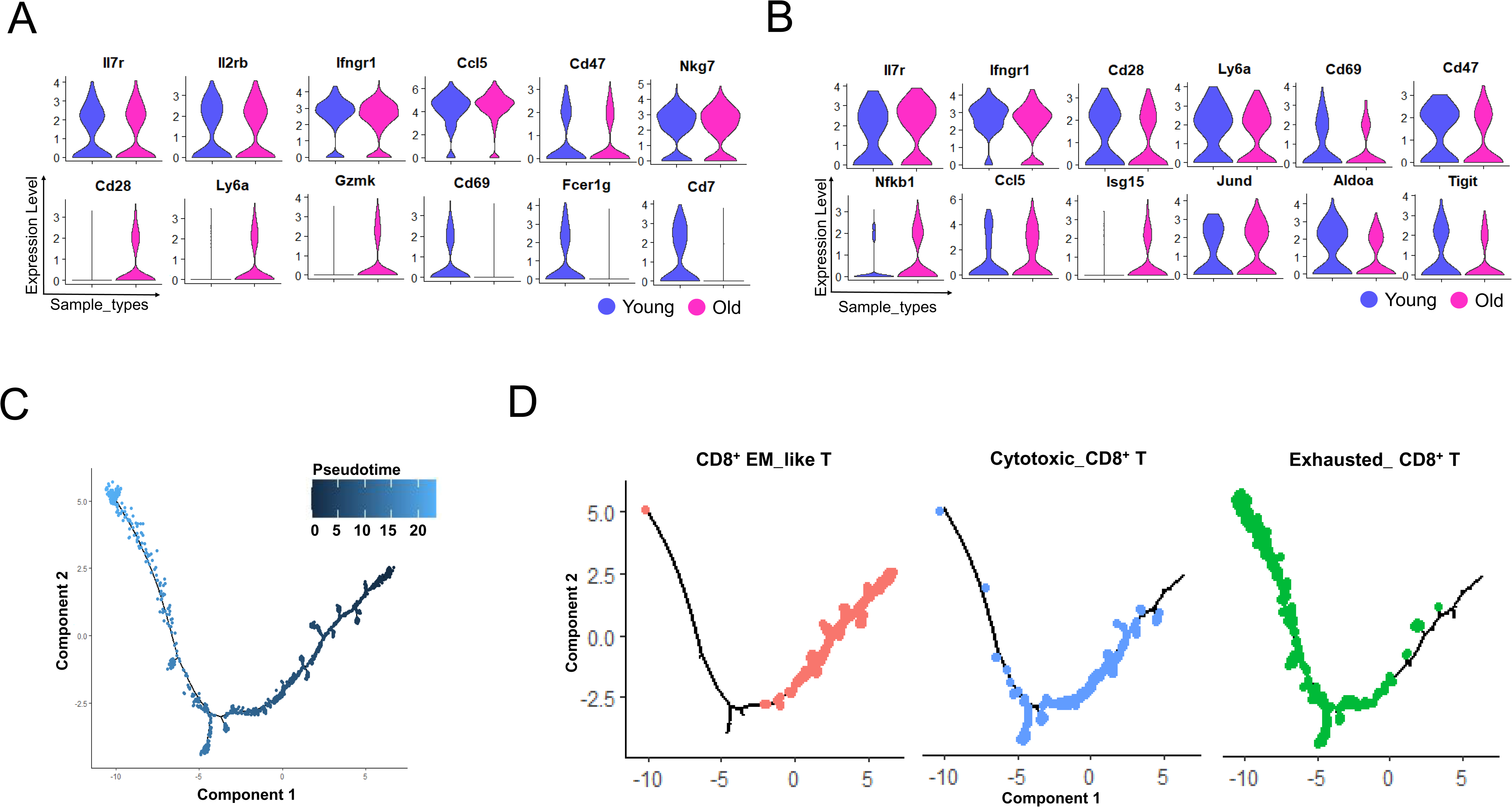
Analysis of effector memory like T cell differentiation status (A and B) Violin plot comparing the expression levels of representative signatures of CD8^+^ EM_like T (A) and CD4^+^ EM_like T (B) in young and old mice. (C) Cell differentiation trajectory of CD8^+^ EM_like T, Cytotoxic CD8+ T and Exhausted CD8^+^ T cells reconstructed by Monocle2 using single-cell RNA-seq data. Color scale indicates the ordering of cells in pseudotime. (D) Cell differentiation trajectory of CD8^+^ EM_like T (left), cytotoxic CD8+ T(middle) and exhausted CD8+ T cells(right) reconstructed by Monocle2 using single-cell RNA-seq data, colored by cell states. The gene expression in A and B was measured as the log_2_ (count + 1).

**Fig. S9.**
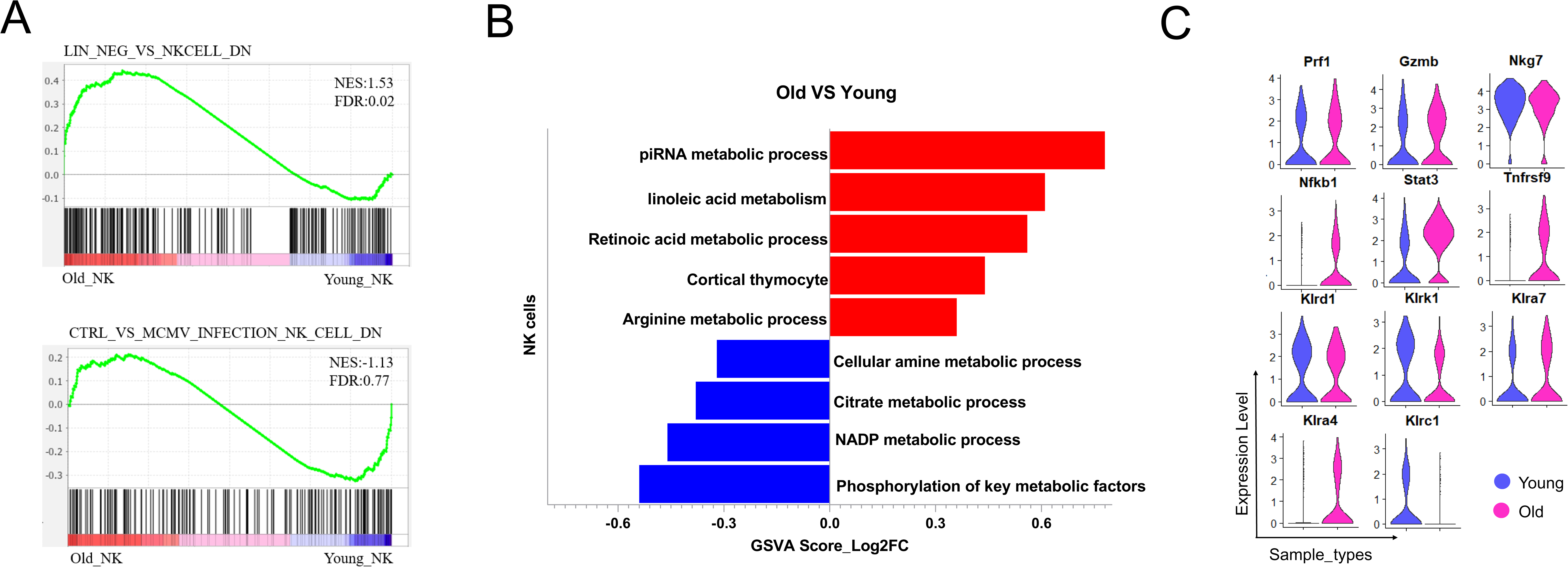
Analysis of tumor-infiltrating NK cells (A) GSEA plots of the signature gene sets in tumor infiltrating NK cells by comparing young with old mice. NES: Normalized Enrichment Score. (B) Gene Set Variation Analysis (GSVA) identified enriched GO terms for NK cells in young vs. old mice. (C) Violin plot showing expression levels of genes associated with NK cell’s function in young and old mice. The gene expression was measured as the log_2_ (count + 1).

**Fig. S10.**
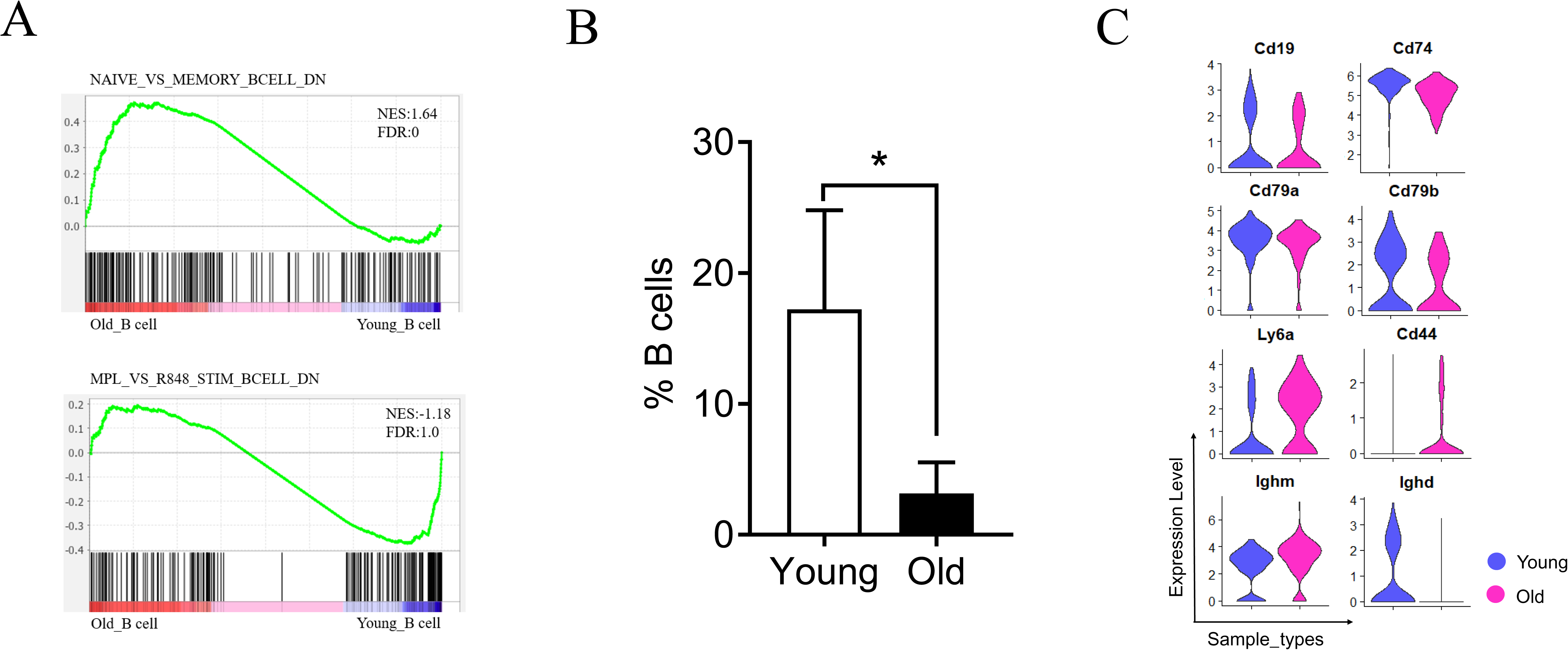
Analysis of tumor infiltrating B cells (A) GSEA plots of the signature gene sets in tumor infiltrating B cells by comparing young with old mice. NES: Normalized Enrichment Score. (B) Proportions of tumor-infiltrating B cells in the young and old mice(n = 3). (C) Violin plot showing expression levels of B cell marker genes in young and old mice. The gene expression was measured as the log_2_ (count + 1). Error bar is mean ± SEM. The significant level is determined by two-tailed, paired Student’s t-test.

